# Single-cell atlas of patient-derived cervical organoids uncovers epithelial immune heterogeneity and intercellular crosstalk during Chlamydia infection

**DOI:** 10.1101/2025.04.13.648603

**Authors:** Pon Ganish Prakash, Naveen Kumar, Stefanie Koster, Christian Wentland, Jayabhuvaneshwari Dhanraj, Rajendra Kumar Gurumurthy, Cindrilla Chumduri

## Abstract

The uterine cervix is a critical mucosal interface that balances immune defense and reproductive function, yet how its distinct epithelial compartments coordinate responses to infection remains unclear. Here, we integrate patient-derived 3D cervical organoids, single-cell transcriptomics and native tissue analysis to construct a high-resolution atlas of epithelial cell diversity and immune dynamics during *Chlamydia trachomatis* infection. We demonstrate that cervical organoids precisely mirror native tissue at both transcriptional and cellular levels, identifying epithelial subtypes with region-specific immune specializations. Upon infection, ectocervical epithelia reinforce barrier integrity, whereas endocervical epithelia, particularly uninfected bystander cells, exhibit extensive transcriptional reprogramming characterized by robust interferon activation, antigen presentation, and antimicrobial defense. Infection profoundly reshapes epithelial intercellular communication, positioning bystander cells as central signaling hubs that coordinate immune responses and tissue regeneration. Our findings highlight a sophisticated epithelial-intrinsic immune network critical for cervical mucosal defense and establish a physiologically relevant platform for studying human host-pathogen interactions and guiding targeted mucosal therapies against reproductive tract infections and pathologies.

## Introduction

Epithelial cells form the frontline of defense across human organ systems, playing pivotal roles in tissue homeostasis, immune surveillance, and pathogen defense ^1,2^. In the female reproductive tract (FRT), the cervical epithelium serves as both a structural and immunological barrier that simultaneously supports reproductive functions and protects against microbial invasion ^3^. The uterine cervix, located at the interface of the upper and lower FRT, comprises two distinct epithelial compartments, the stratified squamous epithelium of the ectocervix and the columnar, mucus-producing epithelium of the endocervix. These regions differ in both function and developmental origin and converge at the squamo-columnar junction (SCJ), a site of active remodeling and high susceptibility to metaplasia and oncogenic transformation^4–6^.

The cervix is a primary entry point for sexually transmitted pathogens, including Chlamydia *trachomatis* (Chlamydia) and high-risk human papillomaviruses (HPVs). Chlamydia, the most prevalent bacterial sexually transmitted infection worldwide, establishes persistent infections that are often asymptomatic yet can lead to serious reproductive complications such as pelvic inflammatory disease (PID), infertility, and ectopic pregnancy ^7,8^. Chronic Chlamydia infection is also implicated in modulating epithelial cell fate, compromising genomic surveillance, and promoting HPV-associated carcinogenesis ^9,10^. Despite the clear relevance of epithelial cell responses in cervical infections, the region- and cell-type-specific immune defense mechanisms of the ecto- and endocervical epithelia remain incompletely defined.

Traditional in vitro models, including immortalized cervical cell lines, have provided foundational insights into Chlamydia pathogenesis but lack the structural, cellular, and immunological complexity of the human cervix ^11,12^. Patient-derived 3D organoid models now offer a powerful alternative. These self-organizing structures preserve tissue-specific lineage fidelity, cellular heterogeneity, and structural polarization, offering unprecedented physiological relevance for infection biology ^13,14^. Recent studies have employed cervical organoids to explore HPV and *Chlamydia* co-infections and epithelial transformation ^15^, yet how distinct epithelial cell types coordinate immune defense and intercellular signaling during infection remains poorly understood.

Single-cell RNA sequencing (scRNA-seq) has revolutionized our ability to dissect epithelial complexity and cellular responses at high resolution. When combined with 3D organoid systems, it enables comprehensive mapping of epithelial differentiation trajectories, infection-induced remodeling, and intercellular communication networks ^16,17^. This integrative approach provides powerful insights into epithelial-intrinsic immunity and the coordinated tissue-level organization of mucosal defense.

In this study, we leverage scRNA-seq and patient-derived ecto- and endocervical organoids to construct a high-resolution cellular atlas of cervical epithelial heterogeneity during Chlamydia infection. We validate the transcriptional fidelity of cervical organoids against native tissue and uncover distinct immune specializations of squamous and columnar epithelia. Upon infection, we observe profound transcriptional reprogramming in endocervical bystander cells, including interferon activation, antimicrobial responses, and activation of antigen-presentation pathways. In contrast, ectocervical cells prioritize structural maintenance and regeneration, exhibiting restrained immune activation. Using ligand-receptor interaction modeling, we show that infection rewires epithelial communication networks, with bystander cells emerging as central hubs for immune coordination and regenerative signaling. Together, our work reveals a compartmentalized yet integrated epithelial defense architecture and establishes a powerful platform for dissecting mucosal immune responses to infection and for designing next-generation precision interventions for FRT diseases.

## Results

### Integrative scRNA-seq analysis of patient-derived cervical organoids and native tissue

To investigate host-pathogen interactions and epithelial cellular dynamics in the cervix, we combined patient-derived cervical organoids with scRNA-seq. Organoids were generated from adult epithelial stem cells isolated from ecto- and endocervical tissues of healthy donors, expanded in lineage-specific cultures, and subsequently infected with GFP-labeled Chlamydia. Infected and uninfected cells were sorted using fluorescence-activated cell sorting (FACS), and high-throughput scRNA-seq was performed using the 10X Genomics platform, capturing epithelial states in both homeostasis and infection (**Fig. 1a**). After quality control, 4,985 high-quality single cells were retained for further analysis.

**Figure 1.**
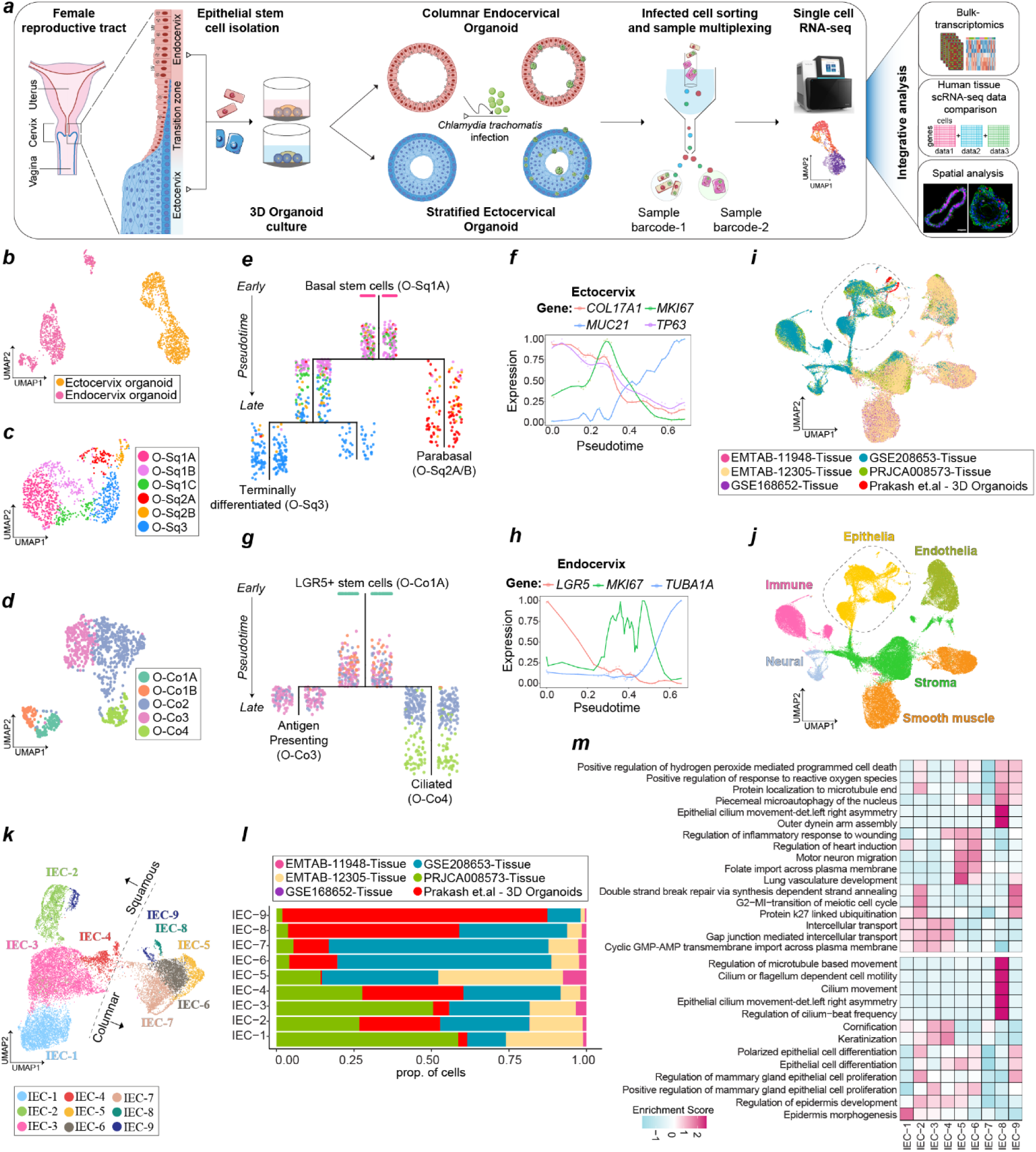
Experimental setup and single-cell characterization of the cervical epithelial landscape using patient-derived 3D organoids. **a)** Schematic of the experimental workflow modeling Chlamydia infection in patient-derived ecto- and endocervical organoids. **b)** Uniform Manifold Approximation and Projection (UMAP) of ecto- and endocervical epithelial cell clusters. Each dot represents a single cell color-coded by tissue type. **c-d)** UMAP projections highlighting squamous (c) and columnar (d) epithelial subclusters, with cells coloured by cluster. **e)** URD differentiation tree of ectocervical squamous epithelial cells; each dot represents a single cell, colored by subcluster. Cells ordered based on pseudotime values starting from early (top) to late (bottom). **f)** Gene expression dynamics of selected squamous markers along the pseudotime; lines represent expression trend of a particular gene. **g)** Differentiation trajectory of endocervical columnar epithelial cells, colored by subcluster and ordered by pseudotime (early to late). **h)** Expression dynamics of key columnar epithelial marker genes along pseudotime. **i)** UMAP visualization of organoid and tissue-derived cell clusters after data integration; cells are color-coded based on their dataset of origin. **j)** UMAP shows six major cell populations across the integrated dataset. **k)** UMAP depicting nine epithelial subclusters identified post-integration, with cells color-coded by cluster. **l)** Bar plot depicting the epithelial cell proportions from different datasets across integrated subclusters. **m)** Heatmap showing gene set enrichment scores for gene ontology (GO) biological processes across integrated epithelial clusters; scale bar denotes the z-scored enrichment values ranging from high (deep pink) to low (blue).

We first examined uninfected organoids to explore epithelial heterogeneity under homeostatic conditions. Unsupervised clustering segregated ecto- and endocervical cells into two transcriptionally distinct clusters, reflecting their lineage specificity and functional divergence (**Fig. 1b**). These findings were further corroborated by microarray analysis (**Fig. S1a-b, Table S1**). Gene ontology (GO) analysis of differentially expressed genes revealed that ectocervical squamous (Sq) epithelial cells were enriched for keratinocyte differentiation and cornification, whereas endocervical columnar (Co) epithelium showed enrichment for cilium assembly and organization (**Fig. S1c**).

Subclustering of the scRNA-seq data revealed six squamous epithelial subtypes (O-Sq1A, O-Sq1B, O-Sq1C, O-Sq2A, O-Sq2B, O-Sq3) and five columnar epithelial subpopulations (O-Co1A, O-Co1B, O-Co2, O-Co3, O-Co4) (**Fig. 1c-d**, **S1d**). Within the squamous lineage, O-Sq1A cells exhibited a stem-like basal phenotype, marked by *COL17A1*, *TP63*, and *ODC1*. O-Sq1B cells were highly proliferative, expressing *CDK1* and *MKI67*, while O-Sq1C cells displayed early differentiation markers, including *KRT14*, and *KRT6A*. Parabasal populations (O-Sq2A, O-Sq2B) expressed *SOX4* and *FOSB*, indicative of transitional states, while terminally differentiated O-Sq3 cells were enriched for *CRNN* and *KRT13*, reflecting epithelial maturation (**Fig. S1e, f-g**).

Pseudotime analysis revealed two distinct differentiation trajectories originating from O-Sq1A basal stem-like cells: one leading to parabasal cells (O-Sq2A, O-Sq2B) and the other toward terminally differentiated squamous cells (O-Sq3) (**Fig. 1e**, **S1h**). As cells progressed along these trajectories, expression of stem cell markers (*COL17A1*, *TP63*) declined, while differentiation markers such as *MUC21* increased, indicating maturation. Notably, *MKI67* expression peaked mid-trajectory, marking a transient proliferative phase before cells reached terminal differentiation (**Fig. 1f**).

In the endocervical epithelium, subclusters O-Co1A and O-Co1B exhibited distinct molecular profiles, with O-Co1A cells expressing *LGR5*, a key stem cell marker essential for glandular epithelial regeneration, whereas O-Co1B cells predominantly expressed *MUC5B* and *MUC5AC*, mucins involved in pathogen defense ^18,19^. Despite their differences, both subclusters expressed *SLC26A2*, *GUCY1A1*, and *DMBT1*, genes associated with epithelial homeostasis and protection against pathogens ^20,21^. O-Co2 cells displayed high *MKI67* and *HMGB2* expression, indicating active proliferation, while O-Co3 cells expressed both Class I (*HLA-A*, *HLA-B*) and Class II (*HLA-DQA1*, *HLA-DRA*) major histocompatibility complex (MHC) genes, suggesting a potential non-professional antigen-presenting cell (np-APC) phenotype. O-Co4 cells, representing highly differentiated ciliated cells, exhibited strong *TUBA1A* and *FOXJ1* expression (**Fig. S1i-k**). Developmental trajectory analysis revealed two differentiation pathways originating from *LGR5*+ O-Co1A stem cells: one leading to np-APCs (O-Co3) and the other to ciliated cells (O-Co4) (**Fig. 1g**, **S1l**). An inverse correlation between *LGR5* and *TUBA1A* expression indicated a progressive shift from stem-like states to terminally differentiated cell types, with *MKI67* expression peaking at the bifurcation point, signifying a proliferative intermediate state (**Fig. 1h**). Differential expression analysis identified gene signatures unique to squamous and columnar subclusters (**Fig. S1f, j, Tables S2-S3**), and cell-type proportion analysis quantified the relative abundance of each epithelial subpopulation within organoid cultures (**Fig. S1m**).

To assess whether patient-derived organoids faithfully recapitulate in vivo transcriptional profiles, we integrated five publicly available human cervical tissue scRNA-seq datasets ^22–26^. Following batch correction, cell clustering was driven by transcriptional identity rather than dataset origin, highlighting robust integration (**Fig. 1i**, **S2a**). Unsupervised clustering identified six major cell types across datasets, including epithelial, stromal, smooth muscle, immune, neural, and endothelial cells (**Fig. 1j**, **S2b**). Notably, epithelial cells from organoids clustered seamlessly with native tissue, confirming their transcriptional fidelity (**Fig. 1i-j**).

Next, re-clustering of the integrated epithelial cell (IEC) population revealed nine transcriptionally distinct subclusters (IEC-1 to IEC-9) (**Fig. 1k**, **S2c**), corresponding to squamous and columnar differentiation states (**Fig. S2d**). IEC-1 to IEC-4 represented squamous epithelial subsets, including basal (IEC-1), proliferative (IEC-2), parabasal (IEC-3), and terminally differentiated (IEC-4) cells. IEC-5 to IEC-9 comprised columnar subtypes, including early progenitors (IEC-5, IEC-6), np-APCs (IEC-7), and ciliated cells (IEC-8). IEC-9, a columnar epithelial population, included a subset of cells that clustered adjacent to proliferative squamous cells (IEC-2), suggesting a shared proliferative transcriptional signature.

We then examined the proportion of epithelial subclusters (IEC-1 to IEC-9) across both organoid and tissue-derived datasets. Notably, patient-derived 3D organoids contributed to all nine epithelial subpopulations, demonstrating their ability to mirror the cellular diversity of native cervical tissue (**Fig. 1l**). To functionally annotate these epithelial subtypes, we performed gene set enrichment analysis (GSEA). Squamous subclusters (IEC-1 to IEC-4) were enriched for epidermis morphogenesis and cornification, while proliferative subclusters (IEC-2, IEC-9) showed enrichment for G2-M cell cycle transition. The ciliated subpopulation (IEC-8) was significantly enriched for cilium motility and beat frequency, confirming its identity (**Fig. 1m**). These findings emphasize the organoid model’s ability to regenerate epithelial heterogeneity and lineage differentiation, making it a reliable system for studying cervical epithelial biology.

### scRNA-seq analysis confirms high transcriptional fidelity of patient-derived cervical organoids to native tissue

To further validate the transcriptional similarity between cervical organoids and native tissue, we extracted squamous and columnar epithelial cells from tissue datasets and performed re-clustering, identifying nine squamous and six columnar subpopulations (**Fig. 2a-b**, **S3a-b**). For clarity and consistency, we uniformly labelled epithelial cells from organoids with ‘O-‘ and from tissue with ‘T-’ prefixes (e.g., O-Co, T-Columnar). Among squamous epithelial subtypes, T-Basal-1 and T-Basal-2 strongly expressed basal markers such as *COL17A1*, *TP63*, and *IGFBP6*, and also expressed *JUN*, *SOX4*, and *FOSB*, genes typically associated with parabasal layers, a pattern not observed in basal squamous cells derived from organoids. T-Cycling-Proliferative-1 and -2 displayed high *MKI67* and *CDK1* expression, indicating active proliferation. T-Parabasal-1 and T-Parabasal-2 were characterized by *KRT6A*, and *MMP7* expression, suggesting an intermediate differentiation state. Terminally differentiated squamous cells, T-Parabasal-Diff-1 to -3, exhibited high expression of *KRT13*, *SPINK5*, and *CRNN*, highlighting late-stage epithelial maturation (**Fig. S3c**).

**Figure 2.**
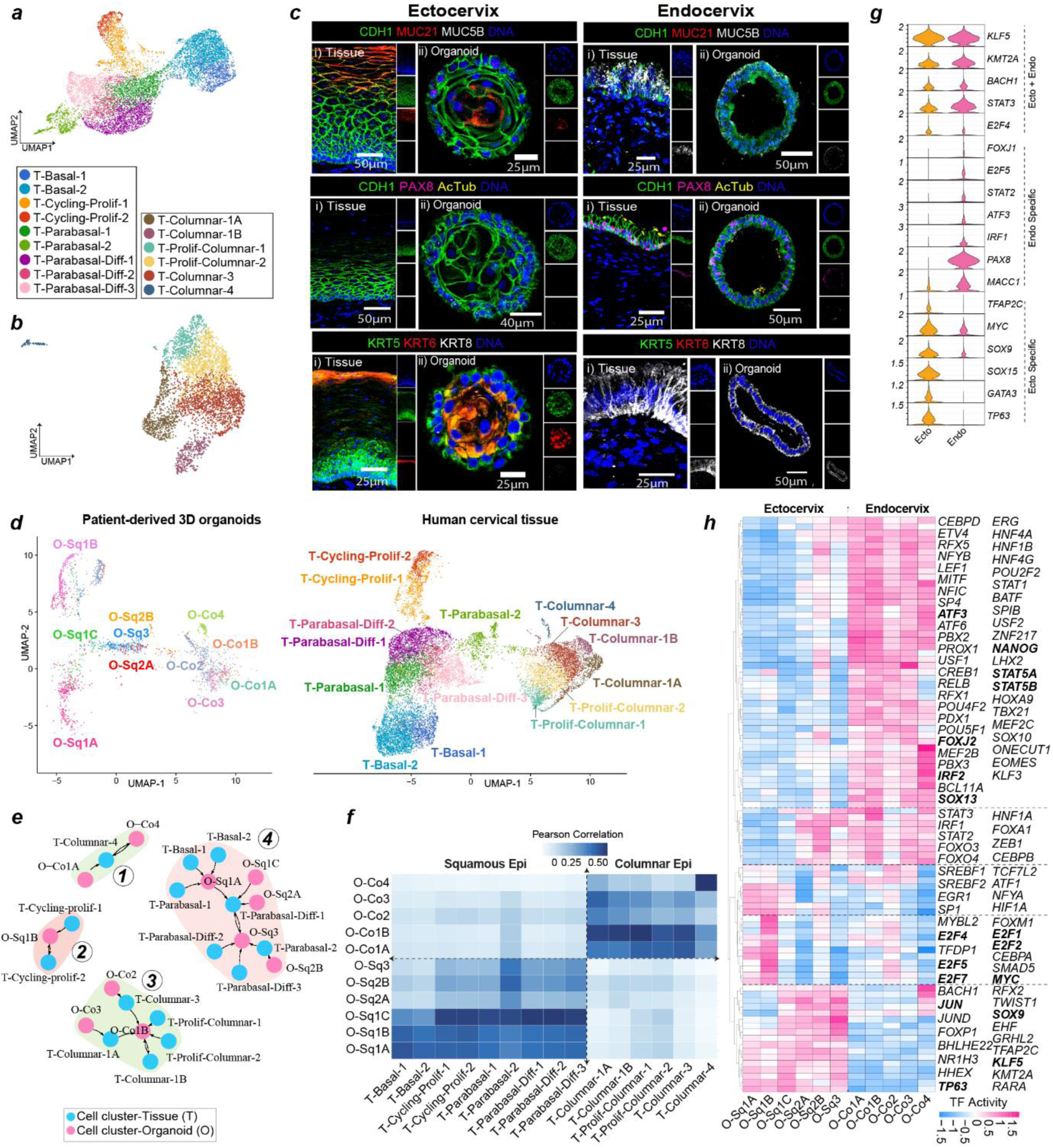
Transcriptional fidelity of patient-derived 3D cervical organoids compared to native cervical tissue. **a-b)** UMAP of tissue-derived squamous (a) and columnar (b) epithelial subclusters; cells colored by cluster annotation. **c)** IHC images of human ectocervix (left panel) and endocervix (right panel) tissues (i) and organoids (ii) stained with CDH1 (green), MUC21(red), MUC5B (gray); and CDH1 (green), PAX8 (magenta), Acetylated tubulin (yellow); KRT5 (green), KRT6 (red), KRT8 (gray). Nuclei were stained with DAPI (blue). **d)** UMAP projection of integrated epithelial cells from organoids (O-) and tissue (T-) datasets, showing annotated subtypes based on transcriptional identity. **e)** Directed graph network showing four distinct communities based on transcriptional similarity between organoid-derived (pink) and tissue-derived (blue) epithelial subclusters; each node represents a cell cluster, and arrows indicate the direction of transcriptional similarity. **f)** Heatmap visualization of Pearson correlation matrix between squamous and columnar epithelial subclusters derived from organoids and tissue. **g)** Violin plot showing normalized gene expression levels of selected transcription factors (TFs) across ecto- and endocervical epithelial subtypes. **h)** Heatmap depicting the most variable TF activity across squamous and columnar epithelial subclusters; color bar indicates activity levels ranging from high (deep pink) to low (blue).

Columnar subtypes displayed a distinct transcriptional landscape. T-Columnar-1A expressed *AXIN2* and *PTGS2*, consistent with an early stem cell-like state, while T-Columnar-1B showed strong expression of *MUC5AC*, *MUC5B*, and *AGR2*, indicating mucus-secreting functions. Proliferative populations, T-Proliferative-Columnar-1 and -2, were enriched for *MKI67* and *PCNA*, whereas T-Columnar-3 expressed MHC genes (*CD74*, *HLA-DPB1*), suggestive of antigen-presenting potential. Ciliated cells (T-Columnar-4) expressed *FOXJ1* and *TUBA1A*, confirming their specialized differentiation state (**Fig. S3d**).

To assess the spatial distribution of epithelial subtypes in tissue and organoids, we performed immunohistochemistry (IHC) (**Fig. 2c**). KRT5 localized to basal and parabasal layers of squamous epithelium, while KRT8 was restricted to columnar cells, consistent with expected lineage identities. KRT6 was enriched in late parabasal and differentiated squamous layers, while MUC21 marked terminally differentiated squamous cells. MUC5B was detected in mucus-secreting columnar cells, while PAX8 and acetylated tubulin (AcTub) confirmed the presence of glandular luminal and ciliated epithelial populations, respectively, in both organoid and tissue samples.

To assess transcriptional fidelity between epithelial subtypes, we first embedded organoid- and tissue-derived epithelial clusters into a shared UMAP space (**Fig. 2d**). This revealed significant overlap, with tight clustering of matched cell types from organoids and tissue, confirming strong transcriptional correspondence. Basal squamous cells from organoids (O-Sq1A) aligned closely with T-Basal-1/2, while proliferative O-Sq1B clustered with T-Cycling-Proliferative-1/2. Parabasal and differentiated squamous populations, including O-Sq1C, O-Sq2A/B, and O-Sq3, mapped to T-Parabasal-1/2, and T-Parabasal-Diff-1/2/3, respectively. Similarly, columnar epithelial subtypes from organoids mapped to their respective tissue counterparts, including O-Co1B with T-Columnar-1B, O-Co2 with T-Columnar-2, and the ciliated O-Co4 population with T-Columnar-4.

To quantify transcriptional similarity between epithelial subtypes of organoid and tissue, we performed cluster similarity analysis by computing similarity scores based on fold-change distributions across shared gene features, ensuring accurate pairing of equivalent cell types across datasets. This analysis revealed four distinct similarity networks, visualized as a community clustering map where nodes represent organoid (O-) or tissue (T-) epithelial populations, and the arrows indicate the direction of transcriptional similarity (**Fig. 2e**). Cluster 1 connected ciliated epithelial cells O-Co4 and T-Columnar-4, indicating strong similarity. Cluster 2 linked proliferative squamous cells across datasets (O-Sq1B and T-Cycling-Proliferative-1/2), while Cluster 3 reinforced the close transcriptional relationship between secretory columnar epithelial cells (O-Co1B and T-Columnar-1B). Cluster 4 grouped basal and parabasal squamous cells, where T-Basal-1/2 and T-Parabasal-1 aligned with O-Sq1A, and O-Sq2B strongly correlated with T-Parabasal-2. Additionally, O-Sq3 exhibited strong similarity to T-Parabasal-Diff-3, emphasizing shared features of terminal differentiation. Pearson correlation analysis of differentially expressed genes further confirms high correlation between organoid- and tissue-derived squamous and columnar populations (**Fig. 2f**).

To investigate lineage-specific transcriptional regulation, we inferred transcription factor (TF) activity from the expression of downstream target genes (**Fig. 2g-h**, **S3e-f**, **Table S4**). Clustering based on TF-activities clearly showed separation between ecto- and endocervical epithelial populations, highlighting distinct regulatory programs underlying their identity and maintenance (**Fig. S3e**). *TP63, JUN, KLF5* and *SOX15* were predominantly active in squamous epithelia ^27,28^, supporting their role in squamous lineage specification. Within squamous subtypes, O-Sq1A and O-Sq1B exhibited high activity of *FOXM1*, *MYC*, and *E2F*, key regulators of cell cycle progression and mitosis ^29,30^. In contrast, columnar epithelial subtypes exhibited strong activity of *PAX8*, *FOXJ1*, *HNF4A, NANOG, STAT5A, RFX5* and *ATF3*, regulators of stem cell fate and ciliogenesis ^31–33^. Shared TFs across both lineages, including *TCF7L2*, *ATF1*, *FOXO3*, and *ZEB1*, suggesting common regulatory pathways.

To further characterize regional epithelial identity, we analyzed HOX gene expression patterns, key regulators of development ^34^. HOXB genes were predominantly expressed in columnar epithelium, while HOXA genes were enriched in squamous subtypes (**Fig. S3g-h**). These patterns reinforce the distinct lineage identities and embryonic origins of ectocervical and endocervical epithelial cells. Together, these findings demonstrate that patient-derived 3D cervical organoids faithfully recapitulate the transcriptional and regulatory landscape of native cervical tissue.

### Differential innate immune architectures of cervical epithelia inform region-specific mucosal defense

Despite the well-established role of epithelia in host defense, the contribution of ectocervical and endocervical epithelial subtypes to innate immune responses remain poorly defined. To address this, we employed microarray and scRNA-seq analysis to profile epithelial innate immune gene expression at both global and single-cell resolution. This approach enabled us to define epithelial subtypes with distinct immune roles, providing a high-resolution view of region-specific immune heterogeneity (**Fig. 3a**).

**Figure 3.**
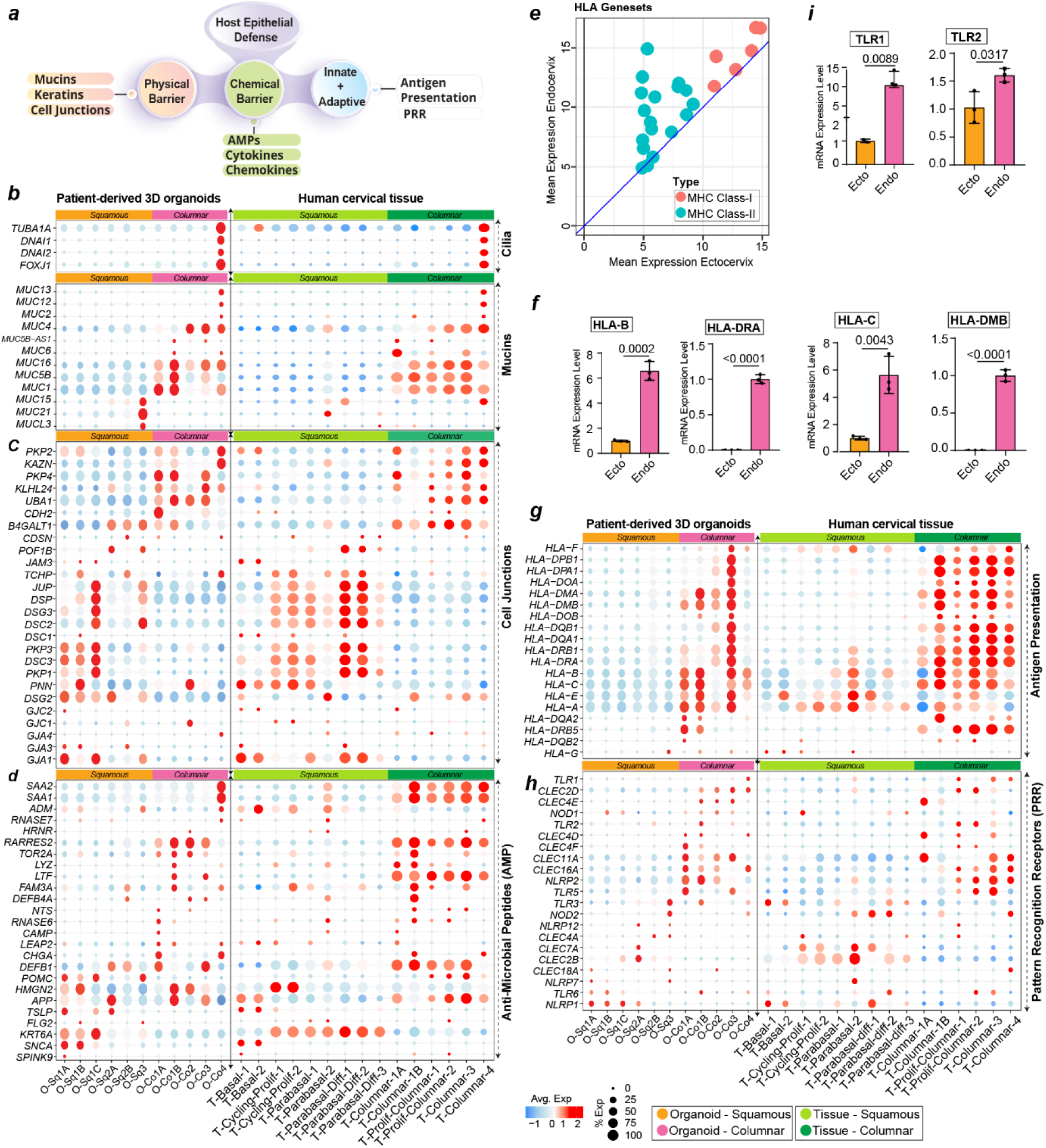
Molecular features of innate immune defense and barrier function in squamous and columnar epithelia of the cervix. **a)** Schematic highlighting the diverse roles of epithelia in host defense and regulation of innate and adaptive immune response. **b-d)** Dot plot showing the expression levels of markers associated with cilia (b), mucins (c) and AMPs (d) in both squamous and columnar epithelial subpopulations from organoids and tissue datasets; Dot size represents the percentage of cells expressing a particular gene; the color bar indicates the intensity of scaled mean expression levels ranging from high (red) to low (blue). **e)** Scatterplot visualization of differentially expressed major histocompatibility complex (MHC) genes between ecto- and endocervical organoids. **f)** qRT-PCR analysis of MHC class I (*HLA-B*, *HLA-C*) and class II (*HLA-DRA*, *HLA-DMB*) gene expression in ecto- and endocervical organoids. **g-h)** Dot plot showing the expression profiles of MHC genes (g) and pattern recognition receptors (PRRs) (h); circle size indicates the percentage of cells expressing a particular gene, and the color bar represents the intensity of scaled mean expression levels from high (red) to low (blue). **i)** qRT-PCR validation of *TLR1*, *TLR2* expression in ecto- and endocervix organoids. Data in (f, i) are presented as mean ± s.d. from three technical replicates, normalized to the ectocervix control, with p-values calculated using Student’s t-test.

To assess barrier integrity across squamous and columnar epithelia, we analyzed the expression of mucins, cytokeratins (KRTs), ciliary genes and intercellular junction markers. Both organoids and tissue exhibited consistent gene expression patterns, validating the reliability of our models. The endocervical columnar epithelium expressed higher levels of both gel-forming and membrane-bound mucins such as *MUC1*, *MUC4*, *MUC5B*, *MUC6*, *MUC12*, *MUC13*, and *MUC16*, compared to the ectocervical cells ^35^. Notably, *MUC2*, *MUC12*, and *MUC13* were enriched in O-Co4 and T-Columnar-4 cells, while *MUCL3*, *MUC15*, and *MUC21* were predominantly expressed in differentiated squamous cells (O-Sq3 and T-Parabasal-diff-3) (**Fig. 3b**, **S4a**). KRTs, critical for mechanical stability and epithelial integrity ^36^, exhibited distinct expression patterns across ectocervical and endocervical subtypes (**Fig. S4b**).

Analysis of cell-cell junctions, including gap, tight, and adherens junctions, revealed distinct organization between ectocervical and endocervical epithelia (**Fig. 3c**, **S4c**). In the ectocervix, *GJA1*, *GJA3*, *DSG2*, *DSC1-3*, *CLDN5*, and *CLDN8* were enriched in basal and parabasal layers, whereas differentiated squamous cells exhibited reduced expression. In contrast, endocervical columnar cells showed robust expression of *GJC1*, *CDH2*, *UBA1*, *PKP4*, and *KAZN*, reflecting a unique structural organization of these cervical epithelia.

A key component of innate host defense is the secretion of antimicrobial peptides (AMPs), which bridge innate and adaptive immunity ^37^. Our analysis of AMP profiles between squamous and columnar epithelial subsets revealed further divergence in immune capacity (**Fig. 3d**, **S4d**). Endocervical columnar cells exhibited strong expression of *DEFB1*, *DEFB4A*, *CHGA*, *LTF*, *LYZ*, *TOR2A*, *HRNR*, and *RARRES2*, suggesting a highly active anti-microbial defense ^38^. Ciliated cells uniquely expressed *SAA1* and *SAA2*, reinforcing their specialized immune surveillance role. Conversely, ectocervical squamous cells displayed limited AMP expression, with *SNCA*, *TSLP*, and *SPINK9* detected, suggesting a greater reliance on physical barrier integrity rather than antimicrobial secretion.

Growing evidence suggests that epithelial cells contribute to antigen presentation, influencing both CD8+ and CD4+ T cell activation ^39,40^. Our analysis revealed that endocervical columnar cells exhibit higher MHC expression compared to ectocervical squamous cells (**Fig. 3e**, **S4e**). qRT-PCR validation confirmed strong MHC expression in endocervical cells, with scRNA-seq identifying O-Co3 cells as np-APCs, exhibiting upregulated MHC class I and II genes (**Fig. 3f-g**). This highlights a previously underappreciated immunomodulatory role of the endocervical epithelium in antigen presentation and mucosal immune priming.

To further characterize pathogen recognition, we examined the expression of pattern recognition receptors (PRRs), including Toll-like receptors (TLRs), nucleotide-binding oligomerization domain-like receptors (NLRs), and retinoic acid-inducible gene-I-like receptors (RLRs) (**Fig. 3h-i**, **S4f**). *TLR1* and *TLR2* expression was significantly higher in columnar epithelial cells, a pattern observed across qRT-PCR, microarray, and scRNA-seq data (**Fig. 3i**). Additional PRRs including *TLR5*, *NLRP2*, *CLEC16A*, and *CLEC4F*, were expressed in columnar subtypes (O-Co1A to O-Co3), reinforcing their role in bacterial recognition and immune activation. These findings align with known ligand specificities, including *TLR2* sensing Gram-positive bacterial components ^41^ and *TLR5* recognizes flagellin, triggering innate immune responses ^42^. Notably, ciliated cells (O-Co4) expressed *TLR1*, *CLEC2D*, and *CLEC4E*, suggesting further immune specialization. In contrast, ectocervical basal squamous cells (O-Sq1A-1C) exhibited high *NLRP1* and *TLR6* expression, while parabasal and differentiated squamous cells expressed *TLR3*, *NOD2*, *CLEC2B*, and *CLEC7A*, indicating distinct PRR-repertoires across cervical epithelial subtypes (**Fig. 3h**).

We next analyzed cytokine and chemokine expression across tissue and organoid (**Fig. S5a-d**), revealing distinct inflammatory and immunomodulatory signatures across ecto- and endocervix. In ectocervical basal squamous cells (O-Sq1A-C), genes linked to inflammatory regulation, including *CCL26*, *IL18*, *IL1RAP*, *IL20RB*, and *IL31RA*, were highly expressed (**Fig. S5d**) ^43^. Parabasal and differentiated squamous cells exhibited high *CXCL14*, *CXCL6*, *CXCL17*, *CXCR2*, *IL1B*, and IL36RN expression. In contrast, endocervical epithelial cells showed strong expression of *CXCL1*, *CXCL3*, *CXCL8*, and *CCL28*, with O-Co4 cells displaying the highest expression. Additionally, O-Co1A cells exhibited *IL23A*, *IL10*, *IL1R1*, *IL1RL2*, *IL18R1*, and *IL1A*, reinforcing their role in immune modulation. While cytokine and chemokine expression were largely consistent between tissues and organoids, some genes displayed differential expression, emphasizing the influential role of tissue microenvironmental cues on epithelial immune regulation.

Together, these findings demonstrate regional specialization of innate immune profiles within the ectocervical and endocervical epithelia, highlighting differences in barrier formation, AMP secretion, antigen presentation, and cytokine signaling. These compartmentalized responses likely contribute to the protection of the FRT against infections.

### Decoding cell type-specific responses of ecto- and endocervical epithelia to Chlamydia infection

To investigate epithelial subtype-specific responses to infections, we evaluated Chlamydia-infected patient-derived ecto- and endocervical organoids (**Fig. 4a**, **S6a**). In ectocervical organoids, Chlamydial inclusions initially appeared in basal layers at 24 hours post-infection (hpi), progressively spreading to adjacent layers by 48 hpi and causing structural disorganization by 4-5 days post-infection (dpi). In contrast, endocervical organoids exhibited increasing GFP fluorescence intensity, indicating intracellular bacterial replication, followed by luminal extrusion of inclusions. Notably, endocervical epithelial cells showed lower infection rates and maintained structural integrity (**Fig. S6a**), suggesting intrinsic differences in susceptibility and response between the two cervical epithelial types.

**Figure 4.**
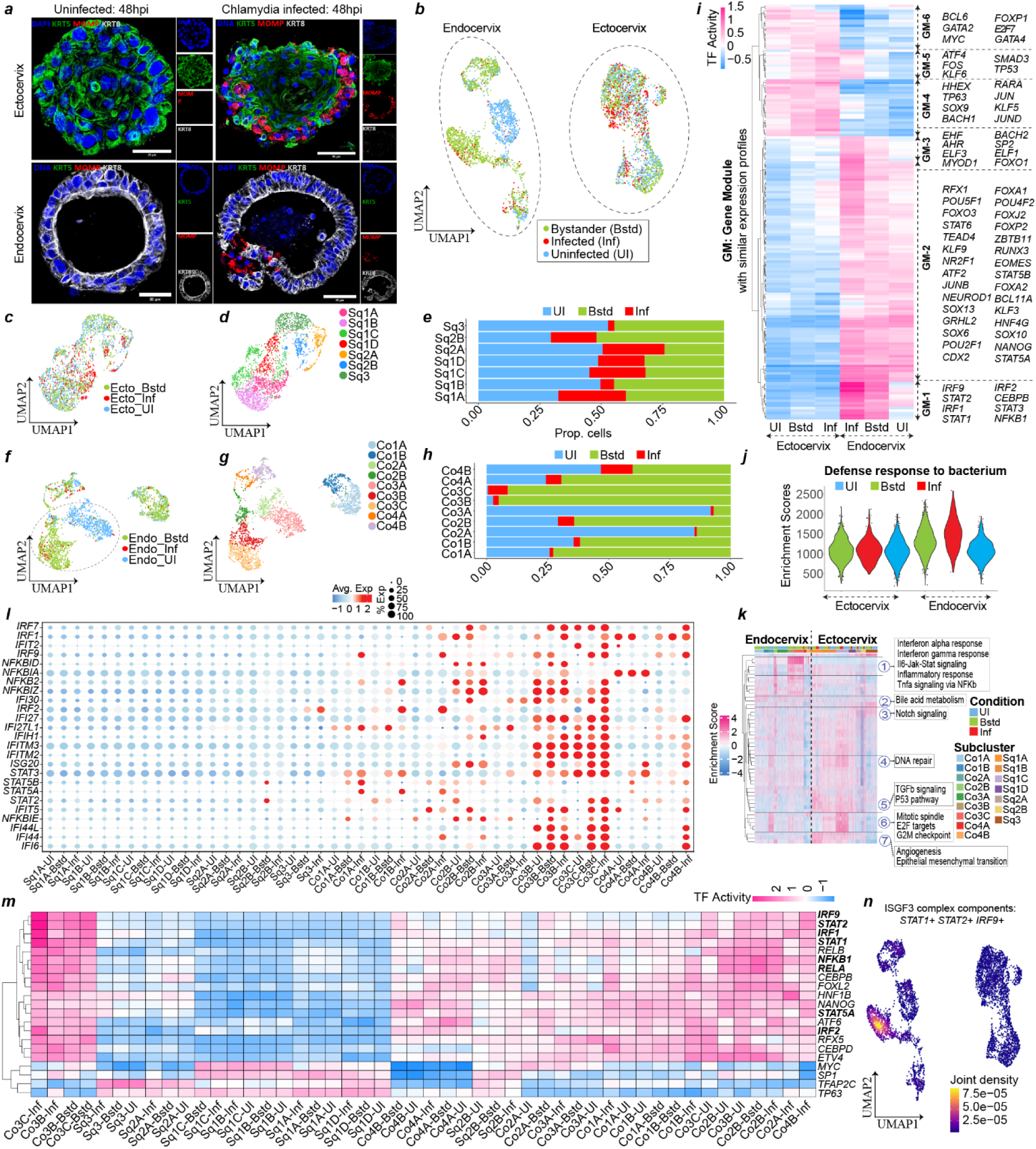
Single-cell transcriptomics reveals cell type-specific responses of ecto- and endocervical epithelia to Chlamydia infection. **a)** IHC images of ectocervix (upper panel) and endocervix (lower panel) organoids, either uninfected (left) or infected (right) with Chlamydia for 48 hours, stained with KRT5 (green), MOMP (red), KRT8 (gray). Nuclei are stained with DAPI (blue). **b)** UMAP projection of uninfected (UI), infected (Inf), and bystander (Bstd) epithelial cells from ecto- and endocervical organoids; each dot represents a single cell, colored by infection status. **c-d)** UMAP visualization of the re-clustered ectocervical squamous epithelial population from (b), colored by infection status (c) and subtypes (d). **e)** Bar plot showing the proportion of UI, Inf, and Bstd cells across squamous epithelial subclusters of the ectocervix. **f-g)** UMAP depicting re-clustered endocervical columnar epithelia from (b), colored by infection status (f) and subtypes (g). **h)** Bar plot shows the proportion of UI, Inf, and Bstd cells across columnar epithelial subclusters of endocervix. **i)** Heatmap of differentially regulated TFs between ecto- and endocervix across infection conditions; color bar depicts the TF activity scores from high (deep pink) to low (blue). **j)** Violin plot showing the gene set enrichment scores for GO term corresponding to defense response to bacterium across epithelial compartments and infection states. **k)** Heatmap showing hallmark pathway enrichment scores across epithelial subtypes and infection conditions. Columns represent individual cells color-coded by tissue and infection status. **l)** Dot plot showing the relative expression of interferon-related genes across ecto- and endocervical subclusters; circle size represents the percentage of cells expressing a particular gene, the color bar indicates the intensity of scaled mean expression levels ranging from high (red) to low (blue). **m)** Heatmap of TFs with highly variable activities across ecto- and endocervical samples and infection states, with subcluster annotations. **n)** Gene-weighted density UMAP projections showing expression of *STAT1*, *STAT2*, and *IRF9* across epithelial cells in (b).

scRNA-seq analysis revealed compartment-specific transcriptional responses to Chlamydia infection (**Fig. 4b**). Within the ectocervical epithelium, seven transcriptionally distinct squamous subclusters (Sq1A-D, Sq2A-B, Sq3) were identified (**Fig. 4c-d**, **S6b-c**). Notably, Chlamydia infected all squamous subtypes, and pseudotime analysis confirmed that differentiation trajectories remained largely unaltered following infection (**Fig. 4e**, **S6b**). In contrast, endocervical epithelia exhibited pronounced transcriptional remodeling, particularly in uninfected bystander cells, which clustered separately from infected cells in UMAP projections (**Fig. 4f**). Nine distinct columnar subpopulations (Co1A-B, Co2A-B, Co3A-C, Co4A-B) were identified (**Fig. 4g**, **S6d-e**), with Co3B and Co3C emerging as transcriptionally remodeled bystander subsets exhibiting unique gene expression signatures (**Fig. S6e**). Pseudotime analysis confirmed that while overall lineage differentiation was preserved, Co3 subsets underwent infection-induced molecular reprogramming. Infections were predominantly localized to the ciliated Co4A and Co4B subpopulations, whereas non-ciliated subsets exhibited relatively low infection rates (**Fig. 4h**). Differential gene expression analysis further delineated infection-induced transcriptional signatures across all epithelial subsets (**Fig. S6f-g**, **Tables S5-6**).

To identify key transcriptional regulators underlying epithelial subtype-specific infection responses, we performed TF activity analysis, revealing five major gene modules (GMs) that distinguished ecto- and endocervical epithelial responses (**Fig. 4i**, **S6j**). GM-1 was characterized by a robust interferon (IFN) response, with enriched activity of *IRF1*, *IRF2*, *IRF9*, *NFKB1*, *STAT1* and *STAT2* in both infected and bystander endocervical cells. This pattern suggests paracrine IFN signaling, wherein infected cells induce interferon-stimulated genes (ISGs) in neighboring epithelial subsets via Type-I and Type-III IFN pathways, a known mechanism of epithelial defense ^44,45^. GM-2 and GM-3 were predominantly associated with columnar epithelial responses and included regulators such as *ATF2*, *CDX2*, *FOXJ2*, and *AHR*, with infection-induced upregulation of TFs such as *SPI1*, *ESRRA*, *RFX1*, and *RARA*. In contrast, GM-4 and GM-5 were primarily associated with squamous epithelial responses, featuring TFs such as *GATA2*, *GATA4*, *SOX9*, and *TP63*, essential for stratified epithelial maintenance. Notably, GM-5 included stress-responsive TFs such as *KLF4*, *ATF4*, *SMAD3*, *FOS*, and *TP53*, which exhibited increased activity across both ectocervical and endocervical subtypes post-infection (**Fig. 4i**, **Table S7**).

To further delineate biological pathways activated in response to Chlamydia infection, we performed enrichment analysis, revealing a clear divergence between squamous and columnar epithelial responses (**Fig. 4j**). The GO term “defense response to bacterium” was significantly enriched in infected and bystander endocervical populations, particularly in Co3B and Co3C subsets. These cells exhibited strong activation of immune pathways, including IFN alpha and gamma responses, IL-6-JAK-STAT signaling, inflammatory response, and TNF-alpha signaling via NF-κB (**Fig. 4k**). In contrast, squamous epithelial populations showed enrichment for Notch and TGF-β signaling, pathways essential for epithelial stratification and barrier homeostasis (**Fig. 4k**, **Table S8**) ^4^. Proliferative subsets, such as Co2A and Sq1C-Sq1D, showed enrichment for E2F target genes and G2/M checkpoint pathways, highlighting an active proliferative response to infection (**Fig. 4k**). Conversely, ciliated endocervical subsets (Co4A-B) showed downregulation of cilium organization and motility-related pathways post-infection, as confirmed by both microarray and scRNA-seq analyses (**Fig. S6h-i**).

At the transcriptional level, columnar subsets, particularly Co3B, Co3C, and infected Co4B, displayed strong upregulation of IFN-stimulated genes (ISGs), including *IFI6*, *IFI27*, *IFITM2*, *IFITM3*, *ISG20*, and *IFI30* (**Fig. 4l**, **S6k-l**). Notably, uninfected bystander cells exhibited comparable ISG expression levels to infected cells, further supporting the hypothesis that paracrine IFN signaling propagates anti-microbial responses throughout the epithelium. In contrast, squamous epithelia showed minimal ISG activation and maintained high *TP63* activity regardless of infection status. Sq1D, Sq2A, and Sq3 subsets exhibited elevated *TFAP2C* activity, while Sq1A-D subsets demonstrated increased activity of *MYC* and *SP1* (**Fig. 4m**, **Table S9**). Columnar subsets exhibited significant upregulation of TFs such as *IRF9*, *STAT2*, *IRF1*, *IRF2*, and *NFKB1*, with the highest activity observed in remodeled populations. Importantly, Co3B and Co3C cells strongly expressed *STAT1*, *STAT2*, and *IRF9* (**Fig. 4n**), suggesting potential assembly of the ISGF3 complex, which translocates to the nucleus to activate ISG transcription.

These findings reveal striking epithelial subtype-specific immune responses to Chlamydia infection. Notably, bystander columnar cells undergo extensive transcriptional reprogramming, mounting a robust pro-inflammatory and IFN-driven response that may serve to limit infection spread and reinforce mucosal immunity.

### Innate immune and anti-microbial responses of cervical epithelial subtypes to Chlamydia infection

To investigate the impact of Chlamydia infection on innate immune responses across epithelial subpopulations, we analyzed gene expression profiles of defense-related markers. Our findings revealed distinct, cell-type-specific immune responses, with both infected and bystander cells showing significant upregulation of innate immunity-associated genes. Notably, epithelial subtypes that expressed mucins and tight junction markers under homeostatic conditions retained their expression post-infection, but at elevated levels in both infected and bystander populations (**Fig. 5a**, **S7a**). For example, differentiated squamous epithelial cells (Sq3) maintained expression of mucins such as *MUC15* and *MUC21*, with increased expression relative to uninfected controls. Similarly, basal and parabasal squamous cells (Sq1 and Sq2) exhibited a pronounced upregulation of PRR genes, including *NLRP1* and *TLR6*. While *NLRP1* plays a key role in microbial defense through inflammasome activation, it has also been implicated in facilitating Chlamydia survival via caspase-1-dependent mechanisms ^46,47^. In contrast, endocervical columnar cells exhibited increased expression of *TLR2*, *TLR5*, and *CLEC4F*, with upregulation observed in both infected and bystander subsets, indicating an active role in bacterial recognition and immune signaling. MHC class I and II gene expression were strongly upregulated in Co1A-B and Co3A-C subpopulations, suggesting enhanced antigen-presenting capacity in columnar epithelial cells, potentially mediated by bystander cell activation (**Fig. 5a**). Consistent with these findings, qRT-PCR confirmed the strong induction of MHC markers and PRR genes in endocervical columnar epithelia, aligning with both microarray and scRNA-seq data (**Fig. 5b-c**, **S7b-c**).

**Figure 5.**
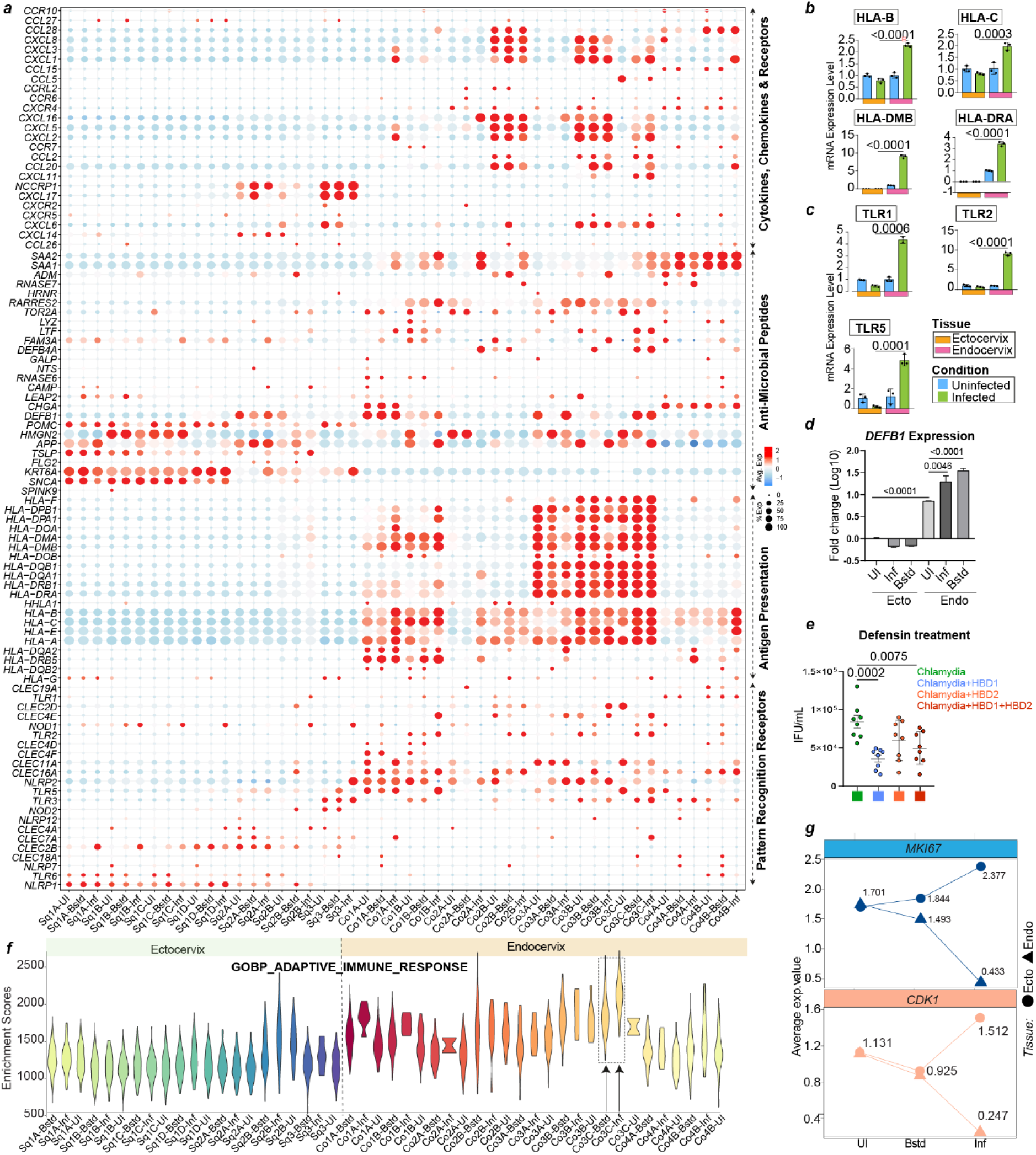
Chlamydia infection alters innate immune and proliferative programs across cervical epithelial subtypes. **a)** Dot plot showing mRNA profiles of PRRs, MHC genes, AMPs and cytokines across ecto- and endocervical epithelial subtypes and UI/Bstd/Inf conditions. The size of each dot reflects the percentage of cells expressing a specific gene, and the color bar signifies the intensity of scaled mean expression levels, ranging from high (red) to low (blue). **b-c)** qRT-PCR analysis of MHC genes (*HLA-B, HLA-C, HLA-DMB, HLA-DRA*) (b); PRRs (*TLR1, TLR2, TLR5*) (c) expression in uninfected and infected ecto- and endocervical organoids at 48hpi. **d)** qRT-PCR analysis of *DEFB1* expression in FACS sorted uninfected, infected, and bystander epithelial cells after 36hpi. Data represent mean ± s.d. from three technical replicates normalized to ectocervix control, and p-values were calculated using Student’s T-test. **e)** Infectivity assay of Chlamydia lysates from ectocervical organoids, with or without pre-treatment with human β-defensins (HBD1, HBD2, or both) at 5 days post-infection (dpi). Data represent mean ± s.d. from 8 non-overlapping regions across two replicates. **f)** Violin plot showing gene set enrichment scores for the GO term related to the adaptive immune response across epithelial subclusters under different infection conditions. **g)** Trend plot showing changes in mean expression levels of selected proliferation markers *MKI67* and *CDK1* under different infection conditions; Lines are colored by gene; point shapes distinguish ectocervical (circles) and endocervical (triangles) samples.

Further analysis of chemical barrier components, including AMPs, cytokines, and chemokines also revealed robust upregulation across infected and bystander cells (**Fig. 5a**, **S7d-e**). Among these, *DEFB1*, showed particularly strong expression in specific endocervical subsets (Co1A-B and Co3A-C) (**Fig. 5a, d**). To assess the functional impact of increased defensin expression, Chlamydia was treated with human β-defensins (HBD1 and HBD2) prior to infection. Chlamydia harvested from cells infected with defensin-treated bacteria exhibited reduced infectivity compared to untreated controls (**Fig. 5e**), indicating that defensins secreted by endocervical epithelial cells can effectively limit Chlamydia infectivity and potentially prevent reinfection.

Pathway enrichment analysis revealed stronger activation of adaptive immune response in endocervical epithelia compared to ectocervical cells, with the highest response observed in Co3C bystander and infected populations (**Fig. 5f**). Among squamous subtypes, Sq2B exhibited the highest immune response scores, suggesting a potential role in coordinating epithelial-immune interactions.

Chlamydia infection has previously been associated with epithelial hyperproliferation ^15^ In line with this, we observed increased expression of proliferation-related markers in infected ectocervical squamous epithelia. However, infected endocervical columnar cells displayed reduced expression of proliferative genes (**Fig. 5g**, **S7f**). These findings suggest that Chlamydia may differentially regulate epithelial proliferation in a cell type-specific manner, possibly influencing tissue remodeling and long-term infection outcomes.

Together, these results highlight the distinct, cell-type-specific innate immune responses of ectocervical and endocervical epithelial subtypes following Chlamydia infection. While squamous epithelial cells prioritize barrier integrity and PRR signaling, columnar cells exhibit a more immune-competent phenotype characterized by enhanced antigen presentation and defensin-mediated antimicrobial defense. These findings underscore the role of epithelial subpopulations in shaping host-pathogen interactions and suggest that the differential immune responses of cervical epithelial compartments may influence infection outcomes and susceptibility to reinfection.

### Coordinated intercellular communication among epithelial subtypes enhances immune and tissue regenerative signaling

Epithelial cell-cell interactions are fundamental for maintaining tissue integrity and coordinating immune responses during infections. Towards this, we used CellChat ^48^ to analyze intercellular signaling networks across homeostatic and Chlamydia-infected states in ecto- and endocervical epithelial subpopulations. We observed a substantial increase in the number of signaling interactions post-infection in both epithelial compartments, suggesting dynamic remodeling of intercellular communication (**Fig. S8a-b**). To dissect these interactions, epithelial subpopulations were categorized as signal senders (cells expressing ligands initiating signaling) and signal receivers (cells expressing cognate receptors that activate downstream pathways). In the ectocervix, signaling pathways such as FGF, CD46, JAM, and DES were consistently enriched under both conditions, whereas IL10, AGRN, and EPGN were exclusive to uninfected states and absent post-infection (**Fig. S8c**). In contrast, pathways such as SEMA4, SEMA6, CADM, and EPHA emerged post-infection, indicating Chlamydia-driven rewiring of epithelial communication (**Fig. S8c, e, Table S10**). Among squamous subtypes, basal Sq1A cells actively engaged in both outgoing and incoming signals involving LAMININ, EGF, and MIF, with signaling strength remaining relatively unchanged after infection (**Fig. S8e**). However, NOTCH signaling exhibited an infection-specific shift, with basal Sq1A cells as primarily signal senders, targeting Sq1C and Sq3 terminally differentiated populations, supporting its role in epithelial stratification ^4^.

Endocervical columnar epithelia displayed distinct infection-induced signaling adaptations. Pathways such as IL1, TNF, NCAM, TRAIL, and GALECTIN were strongly enriched during infection, whereas SEMA4, CDH5, DES, and TGF-β were predominant in uninfected conditions (**Fig. S8d, f, Table S11**). Notably, Co3 cells emerged as key regulators of IL1, TNF, OCLN, and GALECTIN signaling (**Fig. S8f**). IL1 signaling, specifically initiated by Co3 cells, targeted Co1A stem cells, suggesting a potential role in both immune activation and regenerative responses.

To further dissect epithelial communication networks, we examined ligand-receptor (L-R) interactions in squamous and columnar epithelia before and after Chlamydia infection. In squamous epithelial cells, we identified 17 upregulated and 30 downregulated L-R pairs post-infection. Basal (Sq1C) and parabasal (Sq2A-B) cells upregulated ligands such as *AREG*, *LAMB3*, *LAMC2*, and *ANGPTL4*, which are associated with tissue repair and epithelial integrity (**Fig. 6a**) ^49^. Conversely, basal cells exhibited downregulation of antibacterial and differentiation-associated ligands, including *MPZL1*, *BMP7*, *COL4A5*, *DSC2*, and *JAG2* (**Fig. 6b**) ^50,51^. Similarly, parabasal cells exhibited downregulated levels of *DSG2*, *JAG1*, *OCLN*, and *COL4A1*, which are crucial for tight junction formation and pathogen restriction ^52^. suggesting that while epithelial cells activate repair mechanisms, Chlamydia selectively suppresses key host defenses to facilitate its survival.

**Figure 6.**
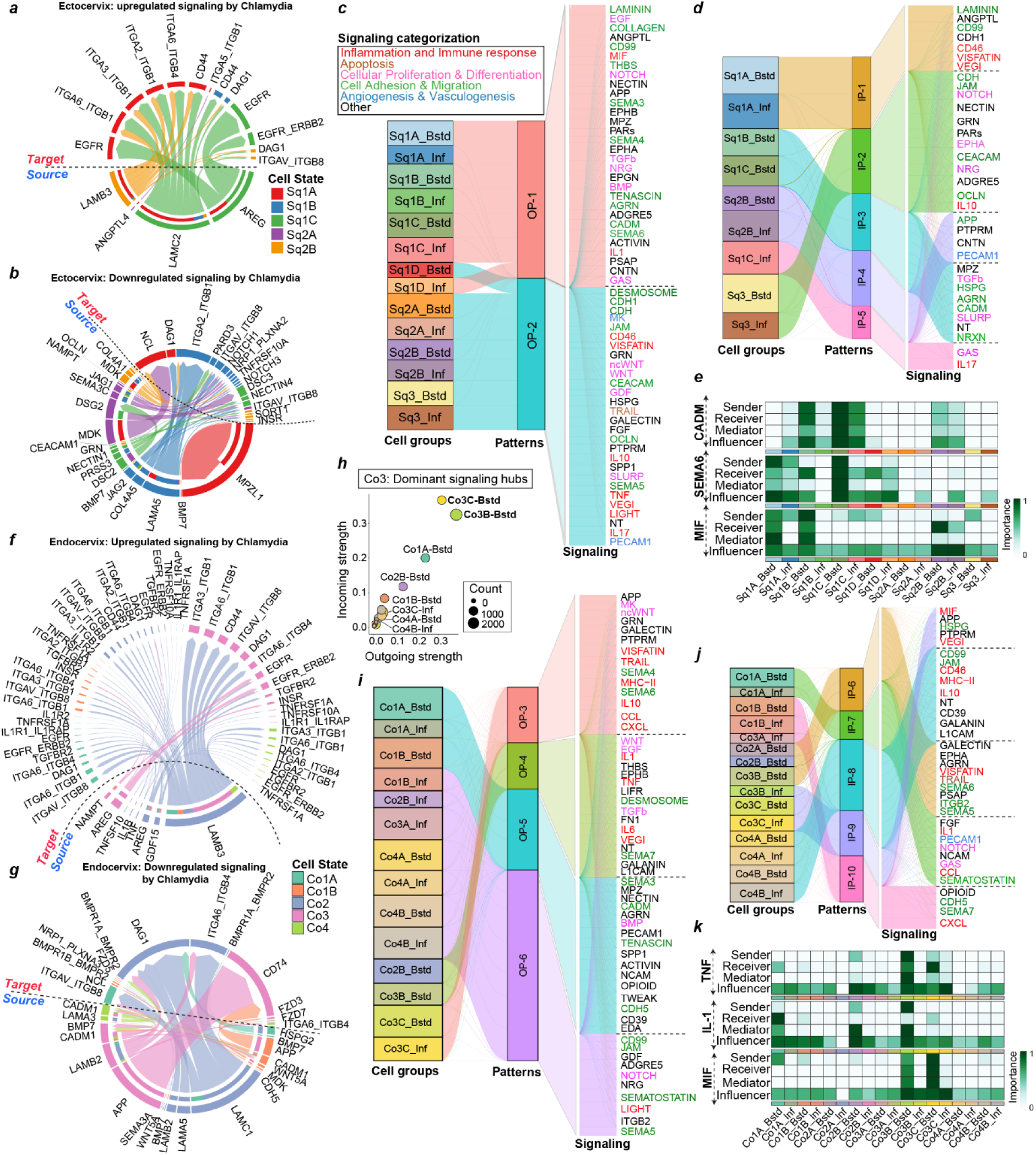
Cell-cell communication analysis reveals differential signaling patterns and networks among ecto- and endocervical epithelial subtypes during Chlamydia infection. **a-b)** Chord diagrams showing upregulated (a) and downregulated (b) ligand-receptor (L-R) signaling pairs in ectocervical squamous epithelial cells upon Chlamydia infection; Outer bars indicate signal sending (ligand-expressing) cell groups; inner bars colored by receiving (receptor-expressing) cell groups; edges colored by signaling source/senders. **c-d)** River plots depicting the inferred outgoing (c) and incoming (d) communication patterns in ectocervical squamous epithelia, highlighting the associations between latent patterns, cell groups, and enriched signaling pathways; thickness of the flow reflects the contribution of cell groups or signaling pathways to each pattern, with pathway labels colored by signaling category. **e)** Heatmap showing the relative importance and signaling role of each cell group in the MIF, SEMA6, and CADM signaling networks in the ectocervix; scale bar color denotes the criticality of a cell group in driving the communication network ranging from high (dark green) to low (gray). **f-g)** Chord diagrams illustrating upregulated (f) and downregulated (g) L–R signaling interactions across endocervical columnar epithelial subpopulations after infection. **h)** Scatter plot visualization of predominant signal senders and receivers among endocervical epithelial subsets in 2D space; Circle size represents the total inferred links (outgoing and incoming) associated with each cell type, colored by subclusters. **i-j)** Inferred outgoing (i) and incoming (j) communication patterns in endocervical epithelia, illustrating the links between latent patterns, cell types, and signaling pathways. **k)** Heatmap visualization of the relative importance and signaling role of each cell group in the endocervix for IL-1, MIF, and TNF signaling networks. Scale bar color as in (e).

Pattern recognition-based signaling analysis in ectocervical epithelia identified two outgoing (OP-1, OP-2) and five incoming (IP-1 to IP-5) signaling patterns (**Fig. 6c-d**). In OP-1, basal Sq1A-C cells were key signal senders, regardless of infection status, actively secreting BMP, TGF-β, SEMA3, and MIF, which regulate immune responses and epithelial regeneration (**Fig. 6c**) ^53,54^. OP-2, predominantly driven by parabasal and terminally differentiated cells, mediated immune and barrier-associated pathways, including CD46, IL17, and JAM. The incoming signaling patterns (IP-1 to IP-5) further emphasized the coordinated response among squamous epithelial subtypes during infection (**Fig. 6d**). For example, IP-1 was specific to basal Sq1A cells, mediating self-coordination via CD99, CD46, and CDH1 signaling. IP-2, targeting terminally differentiated Sq3 cells, involved JAM and NOTCH, reinforcing epithelial barrier integrity, while IP-3 engaged bystander Sq1B and Sq1C cells via TGF-β, AGRN, and SLURP signaling, implicated in tissue remodeling and immune modulation ^53,55^. IP-4, associated with PECAM1, was prominent in parabasal Sq2B cells, while IP-5 was unique to infected, highly proliferative Sq1C cells, targeted by IL17-mediated signaling.

To investigate how infected epithelial cells influence neighboring bystander populations, we mapped directional L-R interactions originating from infected cells and targeting bystander cells (**Fig. S8g**). TNF signaling from Sq2B-infected cells targeted bystander cells expressing TNFRSF1A, while Sq1A and Sq3-infected cells triggered the CD46-JAG1 axis, influencing Sq1A-D bystander cells. The strongest infection-induced interaction involved MIF signaling, where infected epithelial cells expressed MIF, which engage CD74/CD44 receptors on basal (Sq1A-B) and parabasal (Sq2B) bystanders. Network centrality analysis highlighted the critical role of bystander epithelial cells in mediating immune and tissue repair responses, particularly via MIF, SEMA6, and CADM signaling ^56,57^(**Fig. 6e**, **S8g**).

In endocervical columnar epithelia, we identified 15 upregulated and 19 downregulated L-R pairs post-infection. *LAMB3*, *AREG*, *TNF*, and *IL1B* were primarily upregulated in Co2 and Co3 subpopulations, indicating their role as key signal senders (**Fig. 6f**). Conversely, *HSPG2*, which is known to facilitate pathogen entry ^58,59^, was downregulated in Co1A stem-like cells.

Laminin-associated ligands (*LAMC1*, *LAMA5*, and *LAMB2*) and *CADM1*, both implicated in pathogen binding and susceptibility ^60,61^, were also reduced across Co2-Co4 populations (**Fig. 6g**). Remodeled bystander cells of Co3B-3C emerged as central signaling hubs, mediating immune and tissue regenerative signaling, highlighting their pivotal role in coordinating epithelial responses during infection (**Fig. 6h**, **S8f**).

We further identified four outgoing (OP-3 to OP-6) and five incoming (IP-6 to IP-10) signaling patterns in columnar epithelia (**Fig. 6i-j**). Unlike squamous epithelial signaling, columnar epithelial communication was strongly influenced by infection status. Remodeled Co3C cells emerged as key signal senders in OP-3, driving immune-related pathways including IL10, CCL, CXCL, MHC-II, TRAIL, and SEMA4. OP-4 was exclusive to bystander Co2B and Co3B cells, mediating IL1, TNF, TGF-β, and EGF signaling (**Fig. 6i**). Among the incoming signals, IP-6 involved bystander Co2A-B and Co3B populations responding to SEMA5, SEMA6, and GALECTIN signals, while IP-7 linked Co1A stem-like cells to IL1 and CCL-mediated inflammatory signaling (**Fig. 6j**).

Notably, nearly all infected cell types signaled to Co3B and Co3C bystander populations through MIF-(CD74+CD44), a pathway critical for mucosal healing and wound repair ^62^ (**Fig. 6k**, **S8h**). In parallel, TNF signaling from infected Co2B, Co3B, and Co3C cells activated neighboring bystander cells (**Fig. S8h**).

Our study reveals distinct intercellular signaling networks within squamous and columnar epithelia during both homeostasis and Chlamydia infection. While both epithelial lineages engage in signaling pathways that support barrier integrity and tissue repair, endocervical columnar epithelia exhibit a more dynamic and infection-responsive communication profile. Notably, transcriptionally remodeled Co3B and Co3C bystander cells exhibited a unique coordination mechanism, activating a broader set of inflammatory and immune response pathways than their squamous counterparts.

## Discussion

By integrating patient-derived 3D cervical organoids with single-cell transcriptomics and native tissue analysis, our study provides an in-depth, region-resolved high-resolution atlas of cervical epithelial heterogeneity and infection-induced immune dynamics. We show that organoid-derived squamous and columnar epithelial subtypes faithfully recapitulate the molecular, transcriptional, and spatial features of native tissue, supporting the use of these models for interrogating anatomical region-specific host-pathogen interactions. Our findings reveal that stratified ectocervical and columnar endocervical epithelia adopt distinct intrinsic immune strategies to limit infection, preserve tissue integrity, and coordinate responses across cellular compartments (**Fig. 7**).

**Figure 7.**
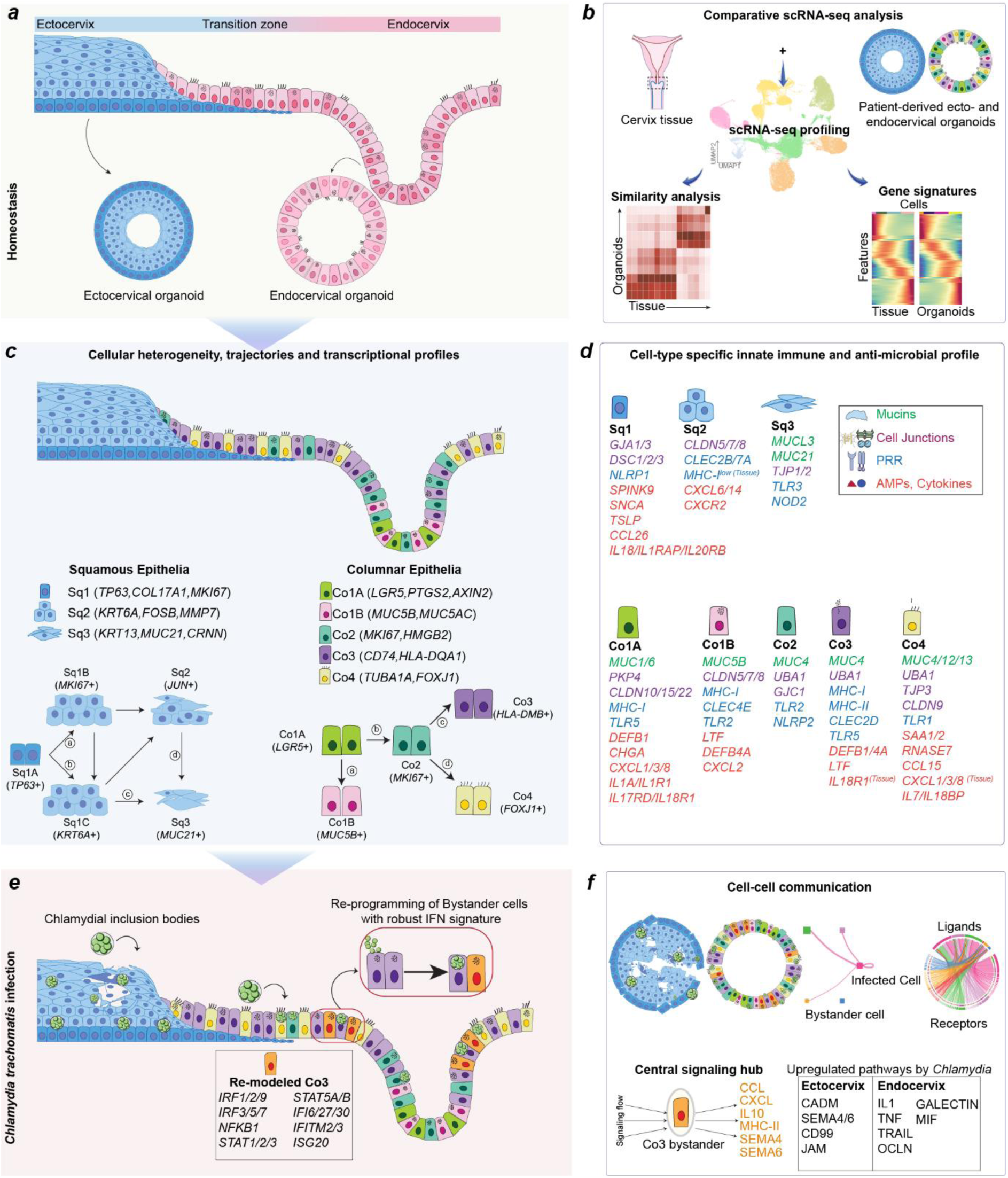
Graphical summary of cervical epithelial cellular diversity, innate mucosal immune features, and Chlamydia-induced responses. **(a)** Anatomical overview of the human uterine cervix highlighting the ectocervix and endocervix, their distinct epithelial compositions, and convergence at the transition zone. Patient-derived 3D organoids derived from adult epithelial stem cells preserve region-specific epithelial architecture and identity. **(b)** Integration of organoid and cervical tissue scRNA-seq datasets reveals high transcriptional fidelity and molecular congruence across corresponding epithelial subsets. Gene signature and similarity analyses demonstrate that organoids recapitulate both cellular heterogeneity and native epithelial programs. **(c)** Depiction shows the identified lineage-specific epithelial subtypes across squamous and columnar compartments under homeostasis. Pseudotime trajectory analysis reveals hierarchical differentiation from basal/stem-like cells to terminally differentiated states, with key subtype-defining transcriptional markers shown. **(d)** Depiction highlighting region- and cell-type-resolved expression profiles of innate immune defense genes at steady state. Distinct epithelial subtypes show selective enrichment of mucins, junctional components, PRRs, antimicrobial peptides (AMPs), and cytokines, reflecting compartmentalized mucosal immune strategies. **(e)** Schematic showing that Chlamydia infection remodels the epithelial landscape. In the endocervix, transcriptional reprogramming of uninfected bystander cells gives rise to IFN-responsive columnar subsets with robust ISG expression, hallmarks of paracrine immune activation. **(f)** Infection reconfigures epithelial communication networks. Cell-cell interaction modeling reveals that columnar bystander cells function as central signaling hubs mediating immune and regenerative responses through pathways such as CCL, CXCL, MHC-II, MIF, SEMA, TNF and IL-1. Region-specific shifts in ligand-receptor interactions underscore divergent mucosal responses between ecto- and endocervix.

Our findings demonstrate that cervical epithelial heterogeneity arises not only from lineage-specific differentiation trajectories but also from regionally distinct immunological specializations. Ectocervical squamous epithelia prioritize and maintain a structural defense strategy enriched in barrier-associated genes and signaling pathways such as NOTCH and TGF-β ^4^. In contrast, endocervical columnar cells displayed pronounced immune competence, characterized by strong baseline and infection-induced expression of mucins, PRRs, cytokines, AMPs, and MHC genes. These findings suggest that endocervical columnar epithelia not only provide physical and chemical defense but also contribute to antigen presentation and immune priming at mucosal surfaces ^39,63^.

Infection with Chlamydia revealed substantial differences in epithelial responses between cervical compartments. In ectocervical organoids, infection was broadly distributed but did not alter differentiation trajectories. The response was characterized by upregulation of stress-responsive transcription factors (e.g., *TP53*, *KLF4*, *ATF4*) and innate sensors (e.g., *NLRP1*, *TLR6*), and a modest increase in PRR and AMP expression while maintaining stratification signatures. This subdued response could reflect a compartmentalized defense strategy optimized for preserving tissue architecture while controlling infection. Notably, one of the most striking findings of this study is the extensive transcriptional remodeling observed in endocervical bystander cells following infection. Despite being uninfected, Co3B and Co3C cells gained a distinct immune-activated state enriched for ISGs, defensins, pro-inflammatory cytokines, and MHC molecules, and exhibited activation of key interferon signaling TFs such as *STAT1*, *STAT2*, and *IRF9*, components of the ISGF3 complex. These features indicate a robust paracrine activation of bystander cells, likely driven by Type-I and Type-III IFNs from infected neighbors, as has been observed in other epithelial systems ^64–66^. Importantly, bystander cells often mounted immune responses comparable to or stronger than infected cells, emphasizing their critical role in amplifying mucosal defense.

This phenomenon of epithelial reprogramming was further reflected in the cell-cell interaction landscape. Chlamydia infection extensively reshaped the ligand-receptor networks, with selective induction and suppression of signaling pathways. In the squamous epithelium of ectocervix, infection promoted CADM, SEMA4, SEMA6, CD99, and JAM signaling, indicative of epithelial remodeling and barrier restoration ^56,67^. NOTCH signaling emerged as a key driver of epithelial stratification post-infection, coordinating basal-to-differentiated transitions and suggesting a repair-oriented program activated in response to epithelial injury. In endocervical epithelia, transcriptionally remodeled Co3B and Co3C bystander subsets emerged as central regulators of infection response. These cells received MIF, TNF, and GAS6 signals from infected neighbors, and, in turn, initiated immunomodulatory signaling via IL10, CXCL, and MHC-II pathways. In parallel, activation of the GAS6-MERTK axis in Co1A stem-like bystander cells points to a regenerative feedback loop regulating inflammation and promoting epithelial repair ^68^. Functionally, we show that epithelial immune gene activation translates to antimicrobial activity. β-defensin (*DEFB1*) expression was robustly induced in endocervical organoids and its exogenous application significantly reduced Chlamydia infectivity when tested in vitro. Given that defensins can also recruit dendritic and T cells ^69^, these findings suggest a dual role in direct pathogen restriction and immune cell recruitment, bridging innate and adaptive immunity.

Together, our findings establish that cervical epithelial subtypes are not passive barriers but are active sensors and regulators of mucosal immunity. They engage in cell-intrinsic defenses, transmit paracrine signals to modulate bystander activation, and coordinate tissue-level communication to limit infection and support regeneration. The compartmentalization of these responses reflects adaptation to the distinct anatomical and functional roles of the ecto- and endocervix. From a translational perspective, these findings offer a framework for identifying immune-competent epithelial subsets (e.g., Co3B/C, Sq2B) and key signaling hubs (e.g., Co3C, Sq1A), as potential targets for modulating mucosal immunity. Interferon signaling and MIF-CD74/CD44 interactions pathways may be leveraged for therapeutic intervention. Moreover, the epithelial remodeling signatures uncovered here may serve as biomarkers of infection-induced dysregulation, with implications for predicting persistent infection, metaplasia, or disease progression to conditions like cancer.

Thus, our study establishes a scalable, region-specific cervical organoid platform integrated with single-cell transcriptomics to dissect host-pathogen interactions. Future extensions incorporating immune cells, microbiota, or co-infections (e.g., HPV) will further elucidate epithelial-immune crosstalk and mucosal homeostasis. Collectively, these findings reveal a highly coordinated, epithelial-intrinsic immune architecture in the human cervix that governs defense and regeneration during infection, informing next-generation mucosal vaccines or epithelial-targeted interventions to protect reproductive health and prevent infection-associated reproductive disorders.

## Materials and Methods

### Antibodies and chemicals

The following primary antibodies were used for immunofluorescence: mouse anti-acetylated tubulin-Alexa-647 (Santa Cruz Biotechnology, sc-23950-AF647), mouse anti-E-cadherin-Alexa-488 (BD Biosciences, 560061), mouse anti-E-cadherin (BD Biosciences, 610181), rabbit anti-KRT5-Alexa488 (Abcam, ab193894 mouse anti-MUC5B (Abcam, ab77995), rabbit anti-MUC21 (ProteinAtlas, HPA052028), rabbit anti-KRT8 (1:200, Abcam, abab59400), mouse-anti-KRT6 (Abcam, ab18586), recombinant rabbit anti-PAX8 (Abcam, ab239363), goat anti-*Chlamydia trachomatis* Major Outer Membrane Protein (MOMP) (1:500, BIO-RAD, 1990-0804) and for labeling the DNA, DAPI (Roche, 10236276001) was used. The following secondary antibodies were used for immunofluorescence: donkey anti-goat Cy3 (1:150, Dianova, 705-165-003), donkey anti-mouse Cy3 (1:150, Dianova, 715-166-020), donkey anti-mouse AlexaFluor-647 (1:150, Dianova, 715-605-150), donkey anti-rabbit AlexaFluor-647 (1:150, Dianova, 711-605-152), goat anti-mouse Cy3 (1:150, Dianova, 115-165-006), goat anti-rabbit Cy3 (1:150, Dianova, 111-165-144) and goat anti-rabbit AlexaFluor-647 (1:150, Dianova, 111-606-045).

### Immunofluorescence histochemistry

Immunofluorescence staining was performed on paraffin-embedded tissue and organoid sections as described previously ^14,15^. Images were acquired with Leica TCS SP5 confocal microscope and processed with Image J.

### Quantitative reverse transcriptase polymerase chain reaction (qRT-PCR) analysis

Total RNA of the uninfected and Chlamydia-infected human ectocervical and endocervical organoid pellets or flow cytometric sorted cell pellets were isolated using AllPrep DNA/RNA Mini Kit (QIAGEN, #80204) according to the manufacturer’s protocol. Using the RevertAid First Strand cDNA Synthesis Kit (Thermo Fisher Scientific, K1621), 1 μg of uninfected and infected human ecto- and endocervix RNA was converted into cDNA according to the manufacturer’s protocol. qRT-PCR was performed using the following reaction mixture: 12.5 ng of cDNA, 3.6 μL of respective forward and reverse primer mix (10 μM) mix, 10 μL of GreenMasterMix (2X) High ROX for qPCR (Genexxon Biosciences, M3052.0000), and made up the volume to 20 μL of DNase/RNase-free water. The following qRT-PCR cycle program was used: 95°C for 10 minutes, followed by 40 cycles at 95°C for 15 seconds and 60°C for 60 seconds. The melting cycle of 95°C for 15 seconds, 60°C for 16 seconds, and 95°C for 15 seconds were used. The expression level of human GAPDH (Glyceraldehyde 3-phosphate dehydrogenase), a housekeeping gene, was used to normalize the relative mRNA levels of respective genes in the uninfected and Chlamydia-infected human ecto- and endocervix organoids. All samples were measured as triplicates. The forward and reverse primers were designed with a melting temperature of 60°C from SigmaAlrich (100 μM) and were diluted to 10 μM before use. The following primer pairs were used: TLR1-forward: 5’-AGTTGTCAGCGATGTGTTCG-3’, reverse: 5’-AAAATCCAAATGCAGGAACG-3’, TLR2-forward: 5’-GGTTCAAGCCCCTTTCTTCT-3’, reverse: 5’-TTCCCACTCTCAGGATTTGC-3’, reverse 5’-TTTCCAGAGCCGTGCTAAGT-3’, TLR5-forward: 5’-TGCCACTGTTGAGTGCAAGTC-3’, reverse: 5’-ACCTGGAGAAGCCGAAGGTAAG-3’, HLA-B-forward: 5’-ATACCTGGAGAACGGGAAGG-3’, reverse: 5’-CAGTGTCCTGAGTTTGGTCCT-3’, HLA-C-forward: 5’-ATTACATCGCCCTGAACGAG-3’, reverse: 5’-GTGTGTCTTTGGGGGTTCTG-3’, HLA-DMB-forward: 5’-CTCACAGCACCTCAACCAAA-3’, reverse: 5’-AGAAGCCCCACACATAGCAG-3’, HLA-DRA-forward: 5’-GAAATGGAAAACCTGTCACCA-3’, reverse: 5’-GCATCAAACTCCCAGTGCTT-3’, CCL2-forward: 5’-CCCCAGTCACCTGCTGTTAT-3’, reverse: GAGTTTGGGTTTGCTTGTCC-3’, DEFB1-forward: 5’-AGATGGCCTCAGGTGGTAAC-3’, reverse: 5’-CACTTGGCCTTCCCTCTGTA-3’.

Relative mRNA levels of the samples were measured in terms of fold-change (2^-(ΔΔCt) by SYBR-Green detection with qRT-PCR, StepOnePlus™ Real-Time PCR System (Applied BiosystemsTM), and StepOneTM Software (v2.3, Applied Biosystems).

### Culturing of organoids and *Chlamydia trachomatis* infection

Human cervical tissue samples were provided by the Department of Gynecology, Charité University Hospital, and August-Viktoria Klinikum, Berlin. Usage for scientific research was approved by their ethics committee (EA1/059/15) and informed consent was obtained from all subjects. The study complies with all relevant ethical regulations regarding research involving human participants. Biopsies were sourced from standard surgical procedures. Only anatomically normal tissue biopsies from anonymous donors were processed within 2-3 h after removal. The ecto- and endocervix organoids were cultured according to the previously described methods ^2,9^. For the infection with the *Chlamydia trachomatis*-L2-GFP strain, organoids were harvested, and Matrigel was removed. For the human endocervix, organoids were broken down before infection by passing the organoid suspension through a 26G needle 4 times. Organoids were infected at MOI of 5 in an infection medium (Advanced DMEM/F12 + HEPES (12mM) + Glutamax (1x) + 5% heat-inactivated fetal calf serum (FCS) for 2 hr at 37°C on a shaker (∼100 rpm). Following infection, organoids were washed, seeded on Matrigel, and allowed to grow in a 3D media for 36 hr for single-cell RNA sequencing and FACS or up to 5 days for bulk transcriptomic analysis and paraffinization.

### AMP treatment and infectivity assay

For AMP treatment, the required amount of bacteria were pretreated with either HBD1 (25 μg/mL, Peptanova, #4337-s) in the presence of 2 mM DTT or HBD2 (25 μg/mL, Peptanova, #4338-s) or in the presence of both in SPG buffer for 30 min at RT. Human endocervical organoids were infected with pretreated bacteria at MOI of 5 for 2 hr in the infection medium. Organoids were washed and allowed to grow in Matrigel for 48 hr. After 48 hr post-infection (hpi), organoids were harvested and centrifuged into pellet. Pellet was transferred into 15 mL falcon tubes with 1mL of infection medium. Approximately 1 mL of sterile glass beads were added to flacon tubes and vortexed for 4 mins. Cells suspension was diluted to 1:2. 50 µL of diluted sample was added to previously grown HeLa cells monolayers in 96 well plates and incubated for 2 hr. Media was replaced with complete media and incubated for 24 hr. After 24hpi, media was removed, added 200 µL of ice-cold methanol incubated plate at 4°C overnight. Methanol was washed twice with 200 µL of 1xPBS. Cells were stained with DAPI (1:1000) in 0.2% BSA solution in PBS for 1 hr. Cells were washed with 1x PBS, images were taken from the fluorescent microscope, and infectivity per mL was calculated.

### Flow cytometric sorting of *C. trachomatis* infected cells

Organoids were incubated with 2 mL of TrypLE in a shaker (20 min, 37°C, 180 rpm for ecto cervix and 10 min, 37°C, 180 rpm for endo cervix organoids). The organoids were made into single cells by pipetting up and down 20 times using a 1 mL pipette tip and diluted with 5 mL of 0.04% BSA-PBS solution. After inverting, cells were passed through a 40 μm cell strainer. The suspension was transferred to 75-mm Falcon tubes with cell strainer caps (Fisher Scientific) and placed on ice. Cells were sorted for GFP positive and negative cells into a Falcon tube containing ice-cold 0.04% BSA-PBS solution using BD FACSAria™ III Cell Sorter (BDbiosciences) running the FACSDiva software (BDbiosciences). Using side-scatter and forward-scatter properties, any cell debris, doublets, or aggregates were excluded. Cells were sorted using GFP (502LP-530/30) filters for positive and negative cells.

### Microarray expression profiling and analysis

Organoids were washed, pelleted, and resuspended in 1 mL Trizol® reagent (Invitrogen™, #15596026). RNA was isolated using the Allprep and RNeasy Mini Kit (Qiagen, #80204) according to the manufacturer’s protocol, and quality was assessed by Agilent 2100 Bioanalyzer with RNA Nano 6000 microfluidics kit (Agilent Technologies, #5067-1511). Microarray experiments were performed as previously described ^15^. Single-color hybridizations were conducted using custom 60K Agilent SurePrint G3 v2 arrays. Raw intensity data were subjected to background correction, quantile normalization, and differential gene expression analysis (with p-Value < 0.05 and 1.5-fold change) of the data were performed using the R package LIMMA ^70^. Over-representation analysis was performed using the function ‘compareCluster’ from the R package ClusterProfiler ^71^ using default settings with significant cutoffs for p-Value and adjusted p-Value (Benjamin-Hochberg-based correction) < 0.05.

### Single-cell RNA seq sample preparation

Organoids from 3 biological replicates of human ectocervical cells and 2 biological replicates of human endocervical cells were harvested (0.04% BSA-PBS) after Chlamydia infection. Single cells were prepared from the organoids as above in 0.04% BSA-PBS solution. After inverting, cells were passed through a 40 μm cell strainer. Single cells were pooled at an equal ratio from biological samples and processed for the CMO labeling using kit 3’ CellPlex Kit Set A (10x Genomics, #PN-1000261). CMO-labeled cells were pooled at an equal ratio, made the cell concentration to 2000 cell/µL, and proceeded for the library preparation. Single cells were partitioned into nanolitre-scale Gel-Bead-In-EMulsions (GEMs) using a 10X Chromium Controller. The library was prepared using Single-Cell 3′ reagent kit v3 for reverse transcription, cDNA amplification, and library construction according to the manufacturer’s protocol. Sequencing was performed in paired-end mode with an S1 100-cycles kit using Illumina Novaseq 6000 sequencer.

### Bioinformatic analysis of scRNA seq data from patient-derived cervical organoids Processing of raw sequencing data and downstream analysis

The CellRanger (v.6.1.1) software from the 10X Genomics platform was used to de-multiplex and process the raw sequencing data of uninfected and Chlamydia-infected ecto- and endocervical organoid samples. We used the ‘cellranger multi’ pipeline with default parameters. Reads were aligned to the 10X reference human genome build GRCh38-2020-A.

Downstream analysis for the generated digital gene expression matrices was performed using the R package Seurat (v.4.1.1) ^72^. Primarily, we scrutinized for potential doublets by neglecting barcodes with less than 100 genes, more than 8000 genes, and more than 80,000 UMI counts. Cells were filtered out if their UMIs derived from the mitochondrial genome exceeded 20%. Additionally, we assessed and excluded the erroneously annotated barcodes, i.e., cells/doublets with a substantial and coherent expression of a hybrid transcriptome based on columnar and squamous epithelial markers such as *LGR5*/*KRT8*/*18* and *TP63*/*KRT5*/*14*, respectively. As an outcome, 4,985 cells were employed for further analyses.

Normalization and variance stabilization of the count data was performed using the R package sctransform (v.0.3.3) ^73^, which also identified the highly variable genes. We regressed out the mitochondrial mapping percentage and cell cycle scores during data normalization and scaling. Dimensionality reduction, clustering, and data visualization were performed using the ‘RunPCA’, ‘FindNeighbors’, ‘FindClusters’, and ‘RunUMAP’ function on the top 30 principal components, with default settings. Lastly, by repeating the same workflow, we re-clustered ecto- and endocervical epithelial cell groups separately to identify the subpopulations present during steady state and infection.

### Differential expression analysis and gene set enrichment analysis (GSEA)

Differentially expressed genes (DEGs) between cell types and clusters were identified using the ‘FindAllMarkers’ function from the Seurat R package, with default parameters. GSEA was performed on scRNA-seq count data using the default functions provided by the R package escape (v.1.4.1) ^74^. The joint gene-weighted density estimation of multiple features was visualized using the R package Nebulosa (v.1.2.0) ^75^.

### Signature scoring analysis

Gene signature scoring for our scRNA-seq data was performed using the R package Ucell (v.1.1.1) ^76^ based on the Mann-Whitney U statistic. Since the scoring technique is rank based, the signature scores are independent of the cellular heterogeneity in the data as they are interpreted as relative expressions of the gene set within an individual cell’s transcriptome. We applied this scoring method to identify the uninfected bystander cells present within the Chlamydia-infected samples, which usually contain both the infected and uninfected bystander cells. Using the purely uninfected and FACS-sorted infected cells, we collected the uninfected and infected cell-specific gene signatures for each cell cluster via differential gene expression analysis. Next, we used the ‘AddModuleScore_UCell’ function to perform gene set scoring in Chlamydia-infected samples. Infected cells had scores higher than defined cutoffs (ranging from the minimum value to the first quartile), whereas the bystander cells secured lower scores and failed to surpass the cutoff value. Later, this information was added back to the scRNA-seq metadata, and the cells from Chlamydia-infected samples were tagged as either bystander or infected cells. Further, we also collected hallmark gene sets of ‘E2F_targets’ and ‘G2M_checkpoint’ for enrichment analysis.

### Cell-Cell Interaction Analysis

We used CellChat’s (v.1.1.3) ^48^ standard pipeline to decode cell-cell communication between squamous and columnar epithelial subsets of uninfected and infected organoids. First, we created individual CellChat objects and preprocessed their expression matrix using the in-built functions ‘identifyOverExpressedGenes’, ‘identifyOverExpressedInteractions’, and ‘projectData’ with default settings to identify potential interactions. We calculated communication probabilities to infer the active signaling networks using the functions ‘computeCommunProb’, ‘computeCommunProbPathway’, and ‘aggregateNet’. In addition, we removed the effect of cell proportion by setting the parameter ‘population.size’ to true during probability calculation. We used the ‘netAnalysis_signallingRole’ function to identify the potential source and targets involved in signaling. Using the ‘identifyCommunicationPatterns’ function, we deduced the coordination between cell types and signaling pathways. Finally, we applied a comparative CellChat framework to comprehensively understand the signaling changes across cell groups pre- and post-infection. Next, we used the function ‘identifyOverExpressedGenes’ to perform differential expression analysis, which revealed the up- and downregulated L-R pairs between the two biological conditions (i.e., uninfected and infected).

### Cell transition trajectory and pseudo-time analysis

Developmental trajectories in our data were modeled using the R package URD (v.1.1.1) ^77^. We used the R package destiny (v. 3.9.1) ^78^ to calculate the transition probabilities between cells and construct the diffusion map. The function ‘plotDimArray’ was used to visualize the diffusion components. Next, we performed pseudotime analysis using the standard in-built functions by defining a set of cells as the starting point (root) and possible endpoint (tip) based on the expression of stem cell and highly differentiated cell type markers, respectively. URD tree structure was generated and visualized using the ‘buildTree’ and ‘plotTree’ functions. Eventually, we modeled the temporal gene regulation using the ‘plotSmoothFit’ function.

### Transcription factor (TF) activity analysis

TF activities in ecto- and endocervical epithelia were investigated using the R package Dorothea (v. 1.4.2) ^79^. Initially, we collected regulons from DoRothEA human regulon database with the highest confidence levels A-C for each TF-target interaction. Here, TF activity is a proxy of the transcriptional state of its direct targets. We calculated the viper scores on Dorothea regulons using the wrapper function ‘run_viper’. Next, we identified the highly variable TFs across cell clusters based on obtained activity scores and transformed them into z-scores for better visualization of results using a heatmap.

### Integrative analysis with cervical tissue scRNA-seq data

To assess transcriptional similarities between in vivo cervical tissue and 3D organoid models, we utilized publicly available scRNA-seq cervical tissue datasets from five sources ^22–26^. Only healthy cervical tissue samples were included for data integration and analysis. For integration with our organoid scRNA-seq data, we employed Seurat v5 ^80^ to address potential batch effects. Each dataset, including our organoid data, was initially processed independently following Seurat’s standard log-normalization workflow. The datasets were subsequently merged into a single Seurat object using the ‘merge’ function, with distinct layers assigned to each dataset via the ‘split’ function. Independent normalization and variable feature detection were then performed across the datasets.

We then performed integrative analysis using the ‘IntegrateLayers’ function with ‘Harmony-based’ integration method. Clustering was performed using the ‘FindNeighbours’ and ‘FindClusters’ functions based on the top 15 principal components, with a resolution of 0.1. Nonlinear dimensionality reduction was performed using the ‘RunUMAP’ function for visualization. As a result, a total of 109,594 integrated cells were obtained. Since the external datasets contained additional cell types such as stromal, immune, endothelial, and smooth muscle cells, we first extracted epithelial cell clusters by identifying epithelial marker genes (*KRT5*, *KRT14*, *KRT13* for squamous epithelia; *KRT7*, *KRT8*, *KRT18*, *PAX8* for columnar epithelia). Re-clustering of epithelial cells from both the organoid and external tissue data was performed using the same integration and clustering workflow. Differential expression analysis was performed by rejoining layers within the Seurat object using the ‘JoinLayers’ function, allowing for subsequent downstream analyses. Cluster similarity between organoid and tissue subpopulations was quantified using the R package ClusterFoldSimilarity with default parameters ^81^. Cell type annotations for identified clusters were performed based on canonical marker gene expression or co-expression, supplemented by information from the Human Protein Atlas database. Further, the clustered gene expression plots were generated using the R package scCustomize ^82^.

## Statistics and reproducibility

GraphPad Prism (v.8) was used for statistical calculations and the generation of plots. The data are displayed as mean ± s.e.m. p<0.05 was considered statistically significant.

## Acknowledgments

We thank the Core Unit SysMed at the University of Würzburg for scRNA-seq. We also thank Shilpa Mary Kurian for assistance with sample preparation. C.C. research is funded by Deutsche Forschungsgemeinschaft (DFG) CH2527/2-1 and The Novo Nordisk Foundation (NNF220C0077183 and NNF23OC0086551). Parts of the project were supported by the German Research Foundation in the CRC 1583 “DECIDE” in projects A08, as well as in the RTG 2157 “3D Infect” in project 12N. The funders had no influence on the study design or analysis of the data.

## Author contributions

C.C. conceived the study; P.G.P., N.K., R.K.G., and C.C. designed the experiments, performed and analyzed the data; N.K, S.K. and J.B.D., performed experimental work and contributed to microarray, IHC and qRT-PCR experiments; N.K. and J.B.D. performed the single-cell preparation and sample multiplexing for scRNA seq. P.G.P performed all the scRNA-seq bioinformatics analysis. C.W. performed microarray bioinformatics analysis with the help of R.K.G and C.C.; P.G.P, R.K.G, and C.C. wrote the manuscript.

## References

1 Marchiando, A. M., Graham, W. V. & Turner, J. R. Epithelial barriers in homeostasis and disease. Annu Rev Pathol 5, 119–144 (2010). 10.1146/annurev.pathol.4.110807.092135

2 Brazil, J. C., Quiros, M., Nusrat, A. & Parkos, C. A. Innate immune cell-epithelial crosstalk during wound repair. J Clin Invest 129, 2983–2993 (2019). 10.1172/JCI124618

3 Gunther, J. & Seyfert, H. M. The first line of defence: insights into mechanisms and relevance of phagocytosis in epithelial cells. Semin Immunopathol 40, 555–565 (2018). 10.1007/s00281-018-0701-1

4 Chumduri, C. et al. Opposing Wnt signals regulate cervical squamocolumnar homeostasis and emergence of metaplasia. Nat Cell Biol 23, 184–197 (2021). 10.1038/s41556-020-00619-0

5 McNairn, A. J. & Guasch, G. Epithelial transition zones: merging microenvironments, niches, and cellular transformation. Eur J Dermatol 21 **Suppl 2**, 21–28 (2011). 10.1684/ejd.2011.1267

6 Giroux, V. & Rustgi, A. K. Metaplasia: tissue injury adaptation and a precursor to the dysplasia-cancer sequence. Nat Rev Cancer 17, 594–604 (2017). 10.1038/nrc.2017.68

7 Witkin, S. S., Minis, E., Athanasiou, A., Leizer, J. & Linhares, I. M. Chlamydia trachomatis: the Persistent Pathogen. Clin Vaccine Immunol 24 (2017). 10.1128/CVI.00203-17

8 den Heijer, C. D. J., et al. Chlamydia trachomatis and the Risk of Pelvic Inflammatory Disease, Ectopic Pregnancy, and Female Infertility: A Retrospective Cohort Study Among Primary Care Patients. Clin Infect Dis 69, 1517–1525 (2019). 10.1093/cid/ciz429

9 Chumduri, C., Gurumurthy, R. K., Zadora, P. K., Mi, Y. & Meyer, T. F. Chlamydia infection promotes host DNA damage and proliferation but impairs the DNA damage response. Cell Host Microbe 13, 746–758 (2013). 10.1016/j.chom.2013.05.010

10 Zadora, P. K. et al. Integrated Phosphoproteome and Transcriptome Analysis Reveals Chlamydia-Induced Epithelial-to-Mesenchymal Transition in Host Cells. Cell Rep 26, 1286–1302 e1288 (2019). 10.1016/j.celrep.2019.01.006

11 Filardo, S., Di Pietro, M. & Sessa, R. Better In Vitro Tools for Exploring Chlamydia trachomatis Pathogenesis. Life (Basel) 12 (2022). 10.3390/life12071065

12 Masters, J. R. HeLa cells 50 years on: the good, the bad and the ugly. Nat Rev Cancer 2, 315–319 (2002). 10.1038/nrc775

13 Chumduri, C. & Turco, M. Y. Organoids of the female reproductive tract. J Mol Med (Berl) 99, 531–553 (2021). 10.1007/s00109-020-02028-0

14 Gurumurthy, R. K., Koster, S., Kumar, N., Meyer, T. F. & Chumduri, C. Patient-derived and mouse endo-ectocervical organoid generation, genetic manipulation and applications to model infection. Nat Protoc 17, 1658–1690 (2022). 10.1038/s41596-022-00695-6

15 Koster, S. et al. Modelling Chlamydia and HPV co-infection in patient-derived ectocervix organoids reveals distinct cellular reprogramming. Nat Commun 13, 1030 (2022). 10.1038/s41467-022-28569-1

16 Triana, S. et al. Single-cell analyses reveal SARS-CoV-2 interference with intrinsic immune response in the human gut. Mol Syst Biol 17, e10232 (2021). 10.15252/msb.202110232

17 Lee, W. et al. A single-cell atlas of in vitro multiculture systems uncovers the in vivo lineage trajectory and cell state in the human lung. Exp Mol Med 55, 1831–1842 (2023). 10.1038/s12276-023-01076-z

18 Seishima, R. et al. Neonatal Wnt-dependent Lgr5 positive stem cells are essential for uterine gland development. Nat Commun 10, 5378 (2019). 10.1038/s41467-019-13363-3

19 Ueda, Y. et al. Cervical MUC5B and MUC5AC are Barriers to Ascending Pathogens During Pregnancy. J Clin Endocrinol Metab 107, 3010–3021 (2022). 10.1210/clinem/dgac545

20 Madsen, J., Mollenhauer, J. & Holmskov, U. Review: Gp-340/DMBT1 in mucosal innate immunity. Innate Immun 16, 160–167 (2010). 10.1177/1753425910368447

21 Kumar, A. et al. A Novel Role of SLC26A3 in the Maintenance of Intestinal Epithelial Barrier Integrity. Gastroenterology 160, 1240–1255 e1243 (2021). 10.1053/j.gastro.2020.11.008

22 Li, C. et al. Single-cell transcriptomics reveals cellular heterogeneity and molecular stratification of cervical cancer. Commun Biol 5, 1208 (2022). 10.1038/s42003-022-04142-w

23 Liu, C. et al. Single-cell dissection of cellular and molecular features underlying human cervical squamous cell carcinoma initiation and progression. Sci Adv 9, eadd8977 (2023). 10.1126/sciadv.add8977

24 Li, C., Guo, L., Li, S. & Hua, K. Single-cell transcriptomics reveals the landscape of intra-tumoral heterogeneity and transcriptional activities of ECs in CC. Mol Ther Nucleic Acids 24, 682–694 (2021). 10.1016/j.omtn.2021.03.017

25 Li, C. & Hua, K. Dissecting the Single-Cell Transcriptome Network of Immune Environment Underlying Cervical Premalignant Lesion, Cervical Cancer and Metastatic Lymph Nodes. Front Immunol 13, 897366 (2022). 10.3389/fimmu.2022.897366

26 Qu, X. et al. Interactions of Indoleamine 2,3-dioxygenase-expressing LAMP3(+) dendritic cells with CD4(+) regulatory T cells and CD8(+) exhausted T cells: synergistically remodeling of the immunosuppressive microenvironment in cervical cancer and therapeutic implications. Cancer Commun (Lond) 43, 1207–1228 (2023). 10.1002/cac2.12486

27 Senoo, M., Pinto, F., Crum, C. P. & McKeon, F. p63 Is essential for the proliferative potential of stem cells in stratified epithelia. Cell 129, 523–536 (2007). 10.1016/j.cell.2007.02.045

28 Sulahian, R. et al. SOX15 governs transcription in human stratified epithelia and a subset of esophageal adenocarcinomas. Cell Mol Gastroenterol Hepatol 1, 598–609 e596 (2015). 10.1016/j.jcmgh.2015.07.009

29 Wang, I. C. et al. Forkhead box M1 regulates the transcriptional network of genes essential for mitotic progression and genes encoding the SCF (Skp2-Cks1) ubiquitin ligase. Mol Cell Biol 25, 10875–10894 (2005). 10.1128/MCB.25.24.10875-10894.2005

30 Muller, H. & Helin, K. The E2F transcription factors: key regulators of cell proliferation. Biochim Biophys Acta 1470, M1–12 (2000). 10.1016/s0304-419x(99)00030-x

31 Colleypriest, B. J. et al. Hnf4alpha is a key gene that can generate columnar metaplasia in oesophageal epithelium. Differentiation 93, 39–49 (2017). 10.1016/j.diff.2016.11.001

32 Lemeille, S. et al. Interplay of RFX transcription factors 1, 2 and 3 in motile ciliogenesis. Nucleic Acids Res 48, 9019–9036 (2020). 10.1093/nar/gkaa625

33 Choksi, S. P., Lauter, G., Swoboda, P. & Roy, S. Switching on cilia: transcriptional networks regulating ciliogenesis. Development 141, 1427–1441 (2014). 10.1242/dev.074666

34 Du, H. & Taylor, H. S. The Role of Hox Genes in Female Reproductive Tract Development, Adult Function, and Fertility. Cold Spring Harb Perspect Med 6, a023002 (2015). 10.1101/cshperspect.a023002

35 Andersch-Bjorkman, Y., Thomsson, K. A., Holmen Larsson, J. M., Ekerhovd, E. & Hansson, G. C. Large scale identification of proteins, mucins, and their O-glycosylation in the endocervical mucus during the menstrual cycle. Mol Cell Proteomics 6, 708–716 (2007). 10.1074/mcp.M600439-MCP200

36 Ramms, L. et al. Keratins as the main component for the mechanical integrity of keratinocytes. Proc Natl Acad Sci U S A 110, 18513–18518 (2013). 10.1073/pnas.1313491110

37 Ma, J. Y., Shao, S. & Wang, G. Antimicrobial peptides: bridging innate and adaptive immunity in the pathogenesis of psoriasis. Chin Med J (Engl) 133, 2966–2975 (2020). 10.1097/CM9.0000000000001240

38 Becknell, B. et al. Expression and antimicrobial function of beta-defensin 1 in the lower urinary tract. PLoS One 8, e77714 (2013). 10.1371/journal.pone.0077714

39 Wira, C. R., Rossoll, R. M. & Young, R. C. Polarized uterine epithelial cells preferentially present antigen at the basolateral surface: role of stromal cells in regulating class II-mediated epithelial cell antigen presentation. J Immunol 175, 1795–1804 (2005). 10.4049/jimmunol.175.3.1795

40 Shenoy, A. T. et al. Antigen presentation by lung epithelial cells directs CD4(+) T(RM) cell function and regulates barrier immunity. Nat Commun 12, 5834 (2021). 10.1038/s41467-021-26045-w

41 Latorre, E., Mendoza, C., Layunta, E., Alcalde, A. I. & Mesonero, J. E. TLR2, TLR3, and TLR4 activation specifically alters the oxidative status of intestinal epithelial cells. Cell Stress Chaperones 19, 289–293 (2014). 10.1007/s12192-013-0461-8

42 Yang, J. & Yan, H. TLR5: beyond the recognition of flagellin. Cell Mol Immunol 14, 1017–1019 (2017). 10.1038/cmi.2017.122

43 Williams, M. A., O’Callaghan, A. & Corr, S. C. IL-33 and IL-18 in Inflammatory Bowel Disease Etiology and Microbial Interactions. Front Immunol 10, 1091 (2019). 10.3389/fimmu.2019.01091

44 Hosey, K. L., Hu, S. & Derbigny, W. A. Role of STAT1 in Chlamydia-Induced Type-1 Interferon Production in Oviduct Epithelial Cells. J Interferon Cytokine Res 35, 901–916 (2015). 10.1089/jir.2015.0013

45 Lazear, H. M., Schoggins, J. W. & Diamond, M. S. Shared and Distinct Functions of Type I and Type III Interferons. Immunity 50, 907–923 (2019). 10.1016/j.immuni.2019.03.025

46 Abdul-Sater, A. A., Koo, E., Hacker, G. & Ojcius, D. M. Inflammasome-dependent caspase-1 activation in cervical epithelial cells stimulates growth of the intracellular pathogen Chlamydia trachomatis. J Biol Chem 284, 26789–26796 (2009). 10.1074/jbc.M109.026823

47 Fenini, G., Karakaya, T., Hennig, P., Di Filippo, M. & Beer, H. D. The NLRP1 Inflammasome in Human Skin and Beyond. Int J Mol Sci 21 (2020). 10.3390/ijms21134788

48 Jin, S. et al. Inference and analysis of cell-cell communication using CellChat. Nat Commun 12, 1088 (2021). 10.1038/s41467-021-21246-9

49 Zaiss, D. M. W., Gause, W. C., Osborne, L. C. & Artis, D. Emerging functions of amphiregulin in orchestrating immunity, inflammation, and tissue repair. Immunity 42, 216–226 (2015). 10.1016/j.immuni.2015.01.020

50 Bao, Z. et al. A potential target gene for the host-directed therapy of mycobacterial infection in murine macrophages. Int J Mol Med 38, 823–833 (2016). 10.3892/ijmm.2016.2675

51 Fang, W. K. et al. Down-regulated desmocollin-2 promotes cell aggressiveness through redistributing adherens junctions and activating beta-catenin signalling in oesophageal squamous cell carcinoma. J Pathol 231, 257–270 (2013). 10.1002/path.4236

52 Van Itallie, C. M., Fanning, A. S., Holmes, J. & Anderson, J. M. Occludin is required for cytokine-induced regulation of tight junction barriers. J Cell Sci 123, 2844–2852 (2010). 10.1242/jcs.065581

53 Worthington, J. J., Fenton, T. M., Czajkowska, B. I., Klementowicz, J. E. & Travis, M. A. Regulation of TGFbeta in the immune system: an emerging role for integrins and dendritic cells. Immunobiology 217, 1259–1265 (2012). 10.1016/j.imbio.2012.06.009

54 Kanth, S. M., Gairhe, S. & Torabi-Parizi, P. The Role of Semaphorins and Their Receptors in Innate Immune Responses and Clinical Diseases of Acute Inflammation. Front Immunol 12, 672441 (2021). 10.3389/fimmu.2021.672441

55 Ertle, C. M. et al. New Pathways for the Skin’s Stress Response: The Cholinergic Neuropeptide SLURP-1 Can Activate Mast Cells and Alter Cytokine Production in Mice. Front Immunol 12, 631881 (2021). 10.3389/fimmu.2021.631881

56 Treps, L., Le Guelte, A. & Gavard, J. Emerging roles of Semaphorins in the regulation of epithelial and endothelial junctions. Tissue Barriers 1, e23272 (2013). 10.4161/tisb.23272

57 Sakurai-Yageta, M., Masuda, M., Tsuboi, Y., Ito, A. & Murakami, Y. Tumor suppressor CADM1 is involved in epithelial cell structure. Biochem Biophys Res Commun 390, 977–982 (2009). 10.1016/j.bbrc.2009.10.088

58 De Pasquale, V., Quiccione, M. S., Tafuri, S., Avallone, L. & Pavone, L. M. Heparan Sulfate Proteoglycans in Viral Infection and Treatment: A Special Focus on SARS-CoV-2. Int J Mol Sci 22 (2021). 10.3390/ijms22126574

59 Zhang, F., Sodroski, C., Cha, H., Li, Q. & Liang, T. J. Infection of Hepatocytes With HCV Increases Cell Surface Levels of Heparan Sulfate Proteoglycans, Uptake of Cholesterol and Lipoprotein, and Virus Entry by Up-regulating SMAD6 and SMAD7. Gastroenterology 152, 257–270 e257 (2017). 10.1053/j.gastro.2016.09.033

60 Manivannan, K., Rowan, A. G., Tanaka, Y., Taylor, G. P. & Bangham, C. R. CADM1/TSLC1 Identifies HTLV-1-Infected Cells and Determines Their Susceptibility to CTL-Mediated Lysis. PLoS Pathog 12, e1005560 (2016). 10.1371/journal.ppat.1005560

61 Kulkarni, A. et al. Oncolytic H-1 parvovirus binds to sialic acid on laminins for cell attachment and entry. Nat Commun 12, 3834 (2021). 10.1038/s41467-021-24034-7

62 Farr, L., Ghosh, S. & Moonah, S. Role of MIF Cytokine/CD74 Receptor Pathway in Protecting Against Injury and Promoting Repair. Front Immunol 11, 1273 (2020). 10.3389/fimmu.2020.01273

63 Wallace, P. K. et al. MHC class II expression and antigen presentation by human endometrial cells. J Steroid Biochem Mol Biol 76, 203–211 (2001). 10.1016/s0960-0760(00)00149-7

64 Stanifer, M. L., Guo, C., Doldan, P. & Boulant, S. Importance of Type I and III Interferons at Respiratory and Intestinal Barrier Surfaces. Front Immunol 11, 608645 (2020). 10.3389/fimmu.2020.608645

65 Vanderheiden, A. et al. Type I and Type III Interferons Restrict SARS-CoV-2 Infection of Human Airway Epithelial Cultures. J Virol 94 (2020). 10.1128/JVI.00985-20

66 He, R., Torres, C. A., Wang, Y., He, C. & Zhong, G. Type-I Interferon Signaling Protects against Chlamydia trachomatis Infection in the Female Lower Genital Tract. Infect Immun 91, e0015323 (2023). 10.1128/iai.00153-23

67 Hagiyama, M. et al. Cell Adhesion Molecule 1 Contributes to Cell Survival in Crowded Epithelial Monolayers. Int J Mol Sci 21 (2020). 10.3390/ijms21114123

68 Nassar, M. et al. GAS6 is a key homeostatic immunological regulator of host-commensal interactions in the oral mucosa. Proc Natl Acad Sci U S A 114, E337–E346 (2017). 10.1073/pnas.1614926114

69 Yang, D. et al. Beta-defensins: linking innate and adaptive immunity through dendritic and T cell CCR6. Science 286, 525–528 (1999). 10.1126/science.286.5439.525

70 Ritchie, M. E. et al. limma powers differential expression analyses for RNA-sequencing and microarray studies. Nucleic Acids Res 43, e47 (2015). 10.1093/nar/gkv007

71 Wu, T. et al. clusterProfiler 4.0: A universal enrichment tool for interpreting omics data. Innovation (Camb) 2, 100141 (2021). 10.1016/j.xinn.2021.100141

72 Hao, Y. et al. Integrated analysis of multimodal single-cell data. Cell 184, 3573–3587 e3529 (2021). 10.1016/j.cell.2021.04.048

73 Hafemeister, C. & Satija, R. Normalization and variance stabilization of single-cell RNA-seq data using regularized negative binomial regression. Genome Biol 20, 296 (2019). 10.1186/s13059-019-1874-1

74 Borcherding, N. et al. Mapping the immune environment in clear cell renal carcinoma by single-cell genomics. Commun Biol 4, 122 (2021). 10.1038/s42003-020-01625-6

75 Alquicira-Hernandez, J. & Powell, J. E. Nebulosa recovers single-cell gene expression signals by kernel density estimation. Bioinformatics 37, 2485–2487 (2021). 10.1093/bioinformatics/btab003

76 Andreatta, M. & Carmona, S. J. UCell: Robust and scalable single-cell gene signature scoring. Comput Struct Biotechnol J 19, 3796–3798 (2021). 10.1016/j.csbj.2021.06.043

77 Siebert, S. et al. Stem cell differentiation trajectories in Hydra resolved at single-cell resolution. Science 365 (2019). 10.1126/science.aav9314

78 Angerer, P. et al. destiny: diffusion maps for large-scale single-cell data in R. Bioinformatics 32, 1241–1243 (2016). 10.1093/bioinformatics/btv715

79 Garcia-Alonso, L., Holland, C. H., Ibrahim, M. M., Turei, D. & Saez-Rodriguez, J. Benchmark and integration of resources for the estimation of human transcription factor activities. Genome Res 29, 1363–1375 (2019). 10.1101/gr.240663.118

80 Hao, Y. et al. Dictionary learning for integrative, multimodal and scalable single-cell analysis. Nat Biotechnol 42, 293–304 (2024). 10.1038/s41587-023-01767-y

81 Gonzalez-Velasco, O. ClusterFoldSimilarity: Calculate similarity of clusters from different single cell samples using foldchanges. (2024). 10.18129/B9.bioc.ClusterFoldSimilarity

82 Marsh, S. E. scCustomize: Custom Visualizations & Functions for Streamlined Analyses of Single Cell Sequencing. 10.5281/zenodo.5706430 (2021).

